# Distinct Bomanins at the *Drosophila 55C* locus function in resistance and resilience to infections

**DOI:** 10.1101/2025.04.16.649162

**Authors:** Yanyan Lou, Bo Zhang, Zhiyuan Zhang, Yingyi Pan, Jianwen Yang, Lu Li, Jianqiong Huang, Zihang Yuan, Samuel Liegeois, Philippe Bulet, Rui Xu, Li Zi, Dominique Ferrandon

**Affiliations:** Sino-French Hoffmann Institute, Guangzhou Medical University, Guangzhou, China; Université de Strasbourg, Strasbourg, France; Modèles Insectes de l’Immunité Innée, UPR 9022 du CNRS, Strasbourg, France; Medical Laboratory of Shenzhen Luohu People’s Hospital, The Third Affiliated Hospital of Shenzhen University, Shenzhen, China; Université Grenoble Alpes, Institute for Advanced Biosciences, Inserm U1209, CNRS UMR 5309, Grenoble, France; Platform BioPark Archamps, Archamps, France; Department of Pediatrics, The Affiliated Foshan Maternal and Children’s Hospital, Guangdong Medical University, Foshan, Guangdong, China

**Author notes:** Equal contributions (Yanyan Lou, Bo Zhang).

## Abstract

Host defense against many Gram-positive bacteria and fungal pathogens is mainly provided by the Toll-dependent systemic immune response in *Drosophila*. While antimicrobial peptides active against these categories of pathogens contribute only modestly to protection, Bomanin peptides are major effectors of the Toll pathway. Remarkably, flies deleted for the *55C* locus that contains 10 *Bomanin* genes are as sensitive as Toll pathway mutant flies to these infections. Yet, the exact functions of single Bomanins in resistance or resilience to infections remain poorly characterized. Here, we have extensively studied the role of these *Bomanin* genes. BomT1 functions in resistance to *Enterococcus faecalis* while playing a role in resilience against *Metarhizium robertsii* infection, like BomS2. BomT1 and BomT2 prevent the dissemination of *Candida albicans* throughout the host, even though they are not sufficient to confer protection to immunodeficient flies against this pathogen in survival experiments. Furthermore, *BomT1* and *BomBc1* mutants are sensitive to an *Aspergillus fumigatus* ribotoxin. We conclude that *55C* Bomanins have defined albeit sometimes overlapping roles in the different facets of host defense against infections.

## Introduction

The major host defense against many Gram-positive bacteria, pathogenic yeasts and molds is mediated by the Toll pathway, which regulates one arm of the systemic humoral immune response (Buchon *et al*, 2014; Ferrandon *et al*, 2007; Lemaitre & Hoffmann, 2007; Lindsay & Wasserman, 2014). This transmembrane receptor is activated by binding to the Spätzle cytokine, which is itself matured into an active Toll ligand by proteolytic cascades triggered upon sensing microbial cell wall components or the catalytic activity of proteases released by invading pathogenic microorganisms (Liegeois & Ferrandon, 2022). Toll in turn activates a specific NF-κB intracellular signaling cascade that ultimately leads to the induction of expression of likely hundreds of genes, including those encoding antimicrobial peptide (AMP) genes (De Gregorio *et al*, 2002; Irving *et al*, 2001; Troha *et al*, 2018). For instance, Drosomycin and Metchnikowin have been shown to have antifungal activity *in vitro* (Fehlbaum *et al*, 1995; Levashina *et al*, 1995). Yet, even though AMPs play a significant role in the host defense against Gram-negative bacteria, they appear to be rather marginally required for protection against Gram-positive bacteria or fungi (Cohen *et al*, 2020; Hanson *et al*, 2019). Remarkably, the deletion of 10 genes of the Bomanin family, members of which had initially been identified by mass-spectrometry analysis of the hemolymph of infected flies (Uttenweiler-Joseph *et al*, 1998), phenocopied the susceptibility of Toll pathway mutants to these categories of pathogens (Clemmons *et al*, 2015). The Bomanin family of 12 genes can be divided in three subgroups: short Bomanins (BomSs) that essentially contain a 16-amino-acid conserved Bomanin domain, tailed Bomanins (BomTs) for which the Bomanin domain is prolonged by a 15 to 82 amino-acid long extension, and bicipital Bomanins (BomBcs) that contain two Bomanin domains separated by a 43 to 103 amino-acid long linker. Functionally, Bomanins are required for a candicidal activity found in the hemolymph of flies and that may require a sufficiently strong expression of BomSs, without much specificity being involved within the distinct BomSs (Lindsay *et al*, 2018). Of note, BomSs but not BomBcs require the activity of a Toll pathway-regulated protein, Bombardier (Bbd), which may be required for the secretion or stability of BomSs (Lin *et al*, 2019).

We have recently reported that an important component of host defense against the opportunistic fungal pathogen *A. fumigatus* is the protection against secreted mycotoxins, which is partially mediated by specific Bomanins (Xu *et al*, 2023), in keeping with another study extending the concept to the defense against secreted virulence factors by another family of Toll pathway effectors, BaramicinA-derived peptides (Huang *et al*, 2023).

While the deletion of subsets of *Bom* genes revealed important information on the function of this family of effector peptides (Clemmons *et al*., 2015), little is known with respect to the function of individual *Bom* genes (Chapman *et al*, 2020; Smith *et al*, 2023; Xu *et al*., 2023). Here, we use complementary genetic strategies to implement the dissection of *55C Bomanin* function in the host defense against a selected set of microbes representing distinct categories of pathogens.

## Results

### The 55C Bomanin locus is required in the host defense against several microbial infections

We first checked that we could reproduce the data published previously when isogenizing the 55C deletions, *Bom^Δ55C^* and *Bom^Δleft^*, in our wild-type genetic background *w* [A5001](Thibault *et al*, 2004) (Fig. 1A). As reported previously, we found that *Bom^Δ55C^* mutant flies were highly susceptible to challenges with *Enterococcus faecalis* and *Candida glabrata* (now called *Nakaseomyces glabratus*). *Bom^Δleft^* mutant flies were either sensitive (*E. faecalis*) or behaved as wild-type flies (*C. glabrata*) as expected (Fig. 1B-C). We next tested additional pathogens. The two *Bom* deficiencies exhibited the same phenotype as *C. glabrata* after the injection of *C. albicans*, even though the two pathogenic yeasts are evolutionarily distant, the latter one being dimorphic (Fig. 1D). Whereas the *Bom^Δ55C^* line was as sensitive as the Toll pathway mutant line *MyD88*, *Bom^Δleft^* displayed an intermediate sensitivity phenotype to an *Aspergillus fumigatus* challenge (Fig. 1E). As reported for the entomopathogenic fungus *Beauveria bassiana*, both *Bom* deficiency lines succumbed rapidly at the same rate to injected spores of a related fungus *Metarhizium robertsii* (Fig. 1F). In contrast, upon a natural infection with the same fungus, *Bom^Δleft^* mutants exhibited an intermediate susceptibility phenotype (Fig. 1G). *Bom^Δ55C^* mutant flies were remarkably at least as sensitive as *MyD88* flies to these infections, in keeping with previous studies (Clemmons *et al*., 2015; Hanson *et al*., 2019; Smith *et al*., 2023).

**Figure 1.**
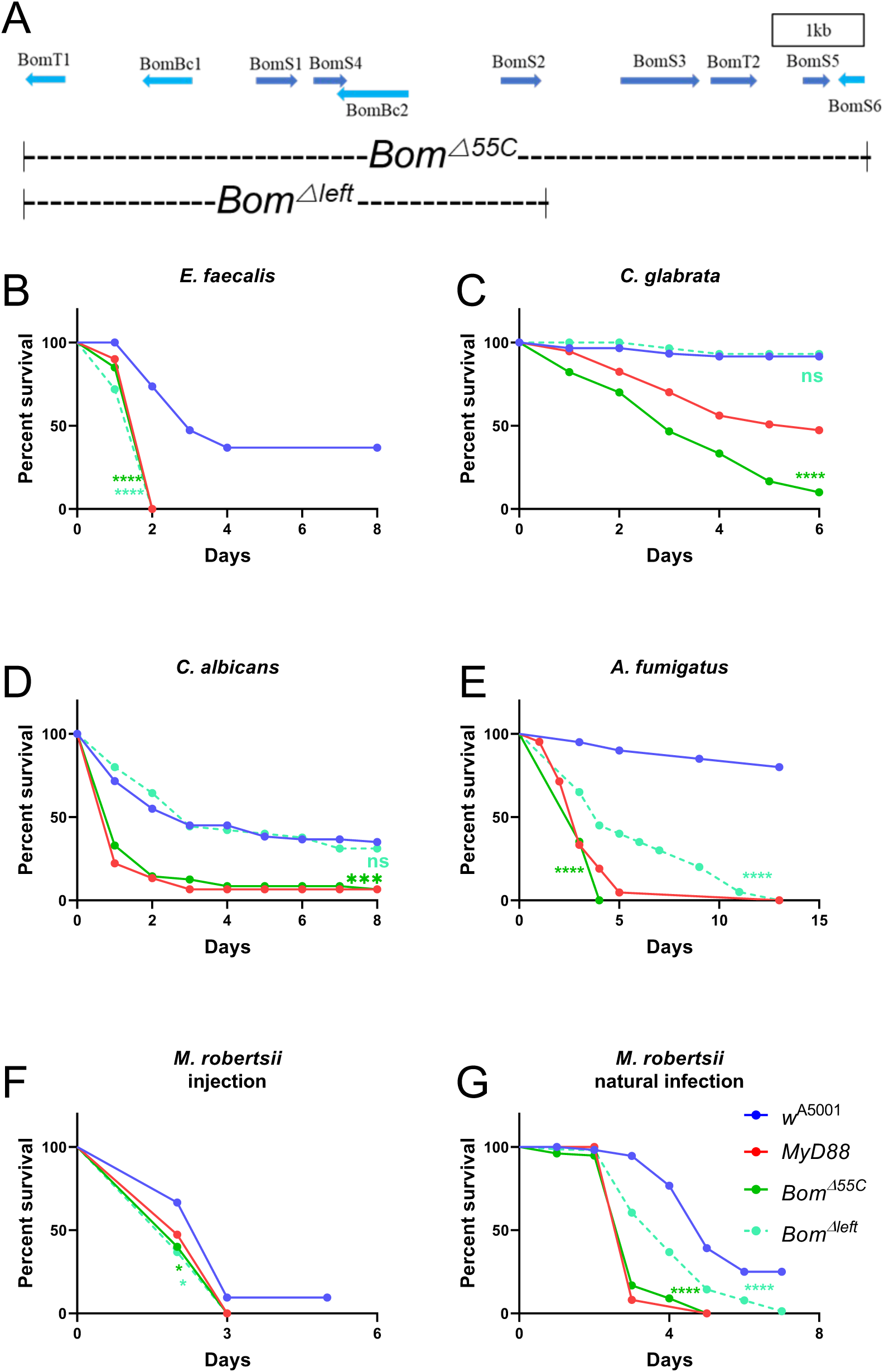
The 55C *Bomanin* locus is essential for host defense against multiple microbial infections. **(A)** The chromosomal localization of the *Bomanin* gene cluster and the deletion intervals of the *Bom^Δ55C^*, *Bom^Δleft^* mutant strains. **(B-G)** Survival curve of wild-type *w* [A5001], *MyD88*, *Bom^Δleft^*and *Bom^Δ55C^* flies with *E. faecalis* **(B)**, *C. glabrata* **(C)**, *C. albicans* **(D)**, *A. fumigatus* **(E)**, *M. robertsii* **(F)** injection and *M. robertsii* **(G)** natural infection. Note: **(B-G)** At least three independent experiments were performed, each experiment used biological triplicates of 20 flies; pooled data are shown. The data were analyzed between infected-mutant and - *w* [A5001] fly using the Log-Rank test; *** p<0.001; **** p<0.0001; ns, no significant difference.

As the host defense encompasses both humoral and cellular immunity, we tested whether *Bom^Δ55C^*mutant flies were defective for the phagocytosis of live *C. glabrata* (Liegeois *et al*, 2020). We did not observe any significant difference in the uptake of these live microorganisms (Fig. EV1).

### Genetic strategies to dissect the different functions of the Bomanins encoded at the 55C locus

The *Bom* genes located at the *55C* locus are required for both resistance and resilience to infections. It is currently not known whether specific *Bomanin* genes are specific to host defense against a given pathogen, whether some are particularly involved in resistance or resilience, whether they may play redundant roles or are required for a “cocktail” effect. Here, we have implemented a dual genetic strategy to address these issues.

First, we have attempted to study the loss-of-function sensitivity phenotype of single *Bom* genes at *55C* after a challenge with either the Gram-positive bacterium *E. faecalis*, the pathogenic yeasts *C. glabrata* and *C. albicans*, the filamentous fungus *A. fumigatus*, or the entomopathogenic fungus *M. robertsii* in two infection models, injection and natural infection. We have generated deletions, knock-out or knock-in mutants using CRISPR-Cas9 technology for *BomT1*, *BomBc1*, *BomS2*, *BomT2*, and *BomS5* (Fig. S1-S6). We expect to have generated null mutants for *BomT1*, *BomBc1*, *BomS2* (two independent mutants), *BomT2* (two deletion lines and one indel), and *BomS5*. Of note, the deletion of *BomS5* severely affects the expression of *BomT2* (Fig. S6). In addition, we have also tested knock-down lines for *BomT1*, *BomBc1*, *BomS4*, *BomBc2*, and *BomT2* by RNA interference (Fig. S7). We have failed to obtain mutant lines affecting the expression of *BomS1*, *BomS3*, and *BomS6*.

Second, we have initiated overexpression strategies to bypass any redundancy issues by overexpressing each gene from transgenes constructed for expression under the control of *UAS* enhancer sequences (Brand & Perrimon, 1993). *Bom* genes were overexpressed first in a wild-type (Fig.S8) and second in a *MyD88* immunodeficient background. However, upon checking the expression of the BomSs in the hemolymph by MALDI-TOF mass-spectrometry (MS)(Uttenweiler-Joseph *et al*., 1998), we failed to observe the expected peaks in *MyD88* flies (MS being semi-quantitative, it was difficult to assess the overexpression of the gene products in the context of an immune response in wild-type flies; in addition, no ectopic expression was detected in the absence of an immune challenge in flies overexpressing *BomS* genes). We reasoned that this effect might be caused by the absence of induction of other *MyD88-*dependent genes that might be required for their expression in the hemolymph. We therefore decided to test each *Bom* gene for a rescuing activity of the sensitivity of *Bom^Δ55C^* deficiency flies to various microbial challenges. Indeed, the role of Bombardier in the secretion or stability of BomS peptide was reported while this work was being pursued (Lin *et al*., 2019). As expected, we then did detect the expression of BomS peptide by MALDI-TOF MS (Fig. S9).

The results we have obtained are summarized in Tables 1 and 2 (a key to understanding Table 1 is provided in Fig. EV2). In the following, we shall describe in more detail the most outstanding results we have obtained.

**Table 1:**
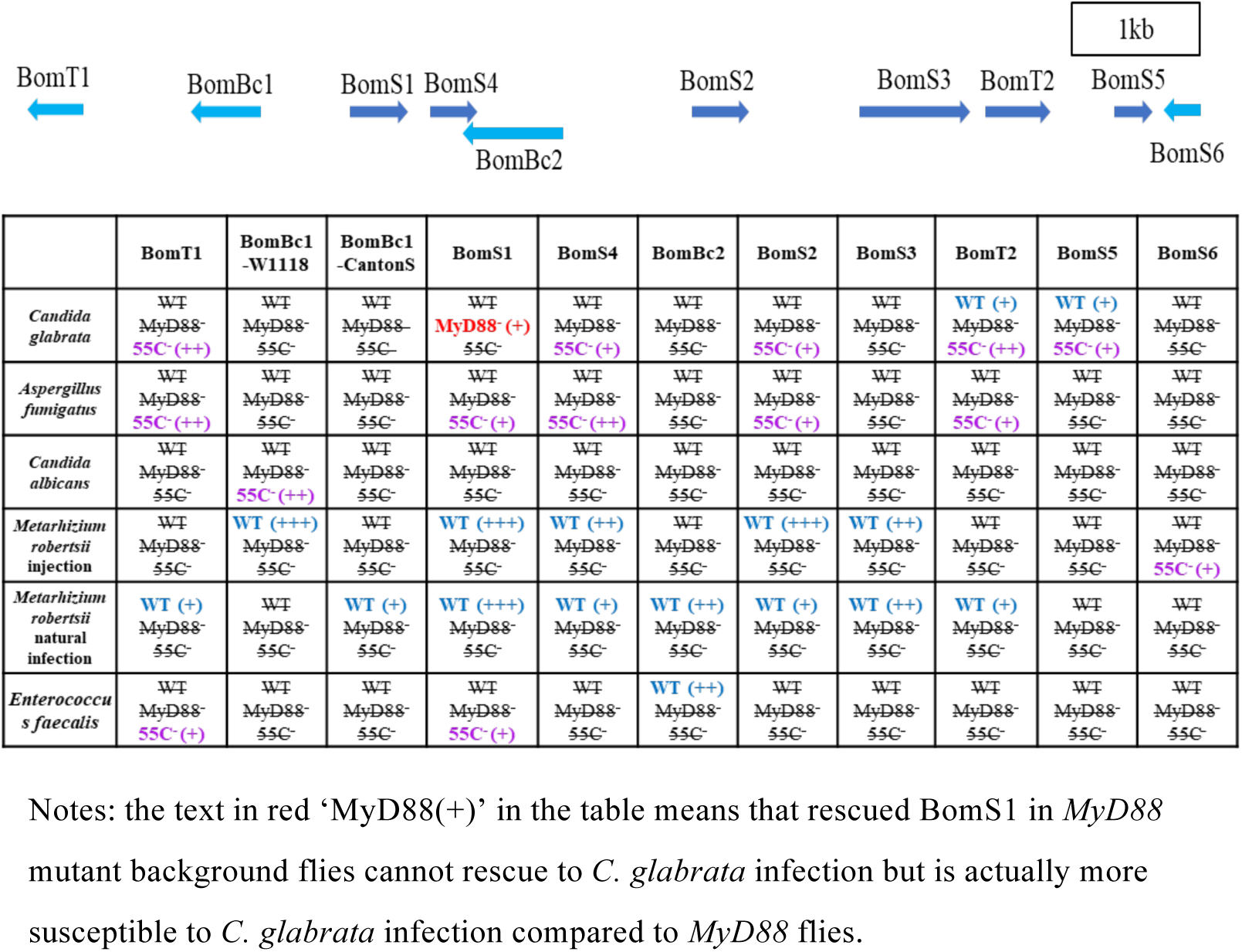
Single overexpressed *Bomanin* genes may improve the protection against specific infections in distinct genetic backgrounds: a summary of *55C* locus *Bomanins*. Notes: the text in red ‘MyD88(+)’ in the table means that rescued *BomS1* in *MyD88* mutant background flies cannot rescue to *C. glabrata* infection but is actually more susceptible to *C. glabrata* infection compared to *MyD88* flies. ‘+’, the intensity of sensitivity. Of note, a key to decipher the indications provided in individual cells is provided in Fig. EV2.

### BomT1 is required but not sufficient for resistance against E. faecalis

When challenged with *E. faecalis*, the *BomT1* knock-in (KI) null mutant flies succumbed at a rate that was almost as fast as *MyD88* mutants (Fig. 2A). A similar, albeit milder, phenotype was observed when *BomT1* was silenced ubiquitously only at the adult stage (Fig. 2B). Of note, the overexpression of *BomT1* in wild-type flies did not protect them from *E. faecalis* (Table 1, Fig. S10A); however, it conferred a partial protection to *Bom^Δ55C^* deletion mutants, suggesting it may need to act in concert with another *55C Bomanin* gene for protection to wild-type levels (Fig. 2C). We have so far failed to identify such a gene in our loss-of-function approach, even though one of the two null *BomS2* mutant line displayed a sensitivity to *E. faecalis* (Table 1, Fig. EV3A). We also noted that *BomS1* overexpression also partially rescued the *Bom^Δ55C^* deletion sensitivity phenotype (Table 1, Fig. EV3B). Unexpectedly, *BomBc2* overexpression provided a degree of protection against *E. faecalis* in wild-type but not *Bom^Δ55C^* flies (Table 1, Fig. EV3C), suggesting it may also act in concert with another 55C *Bom* gene, *BomT1* being the best candidate.

**Figure 2.**
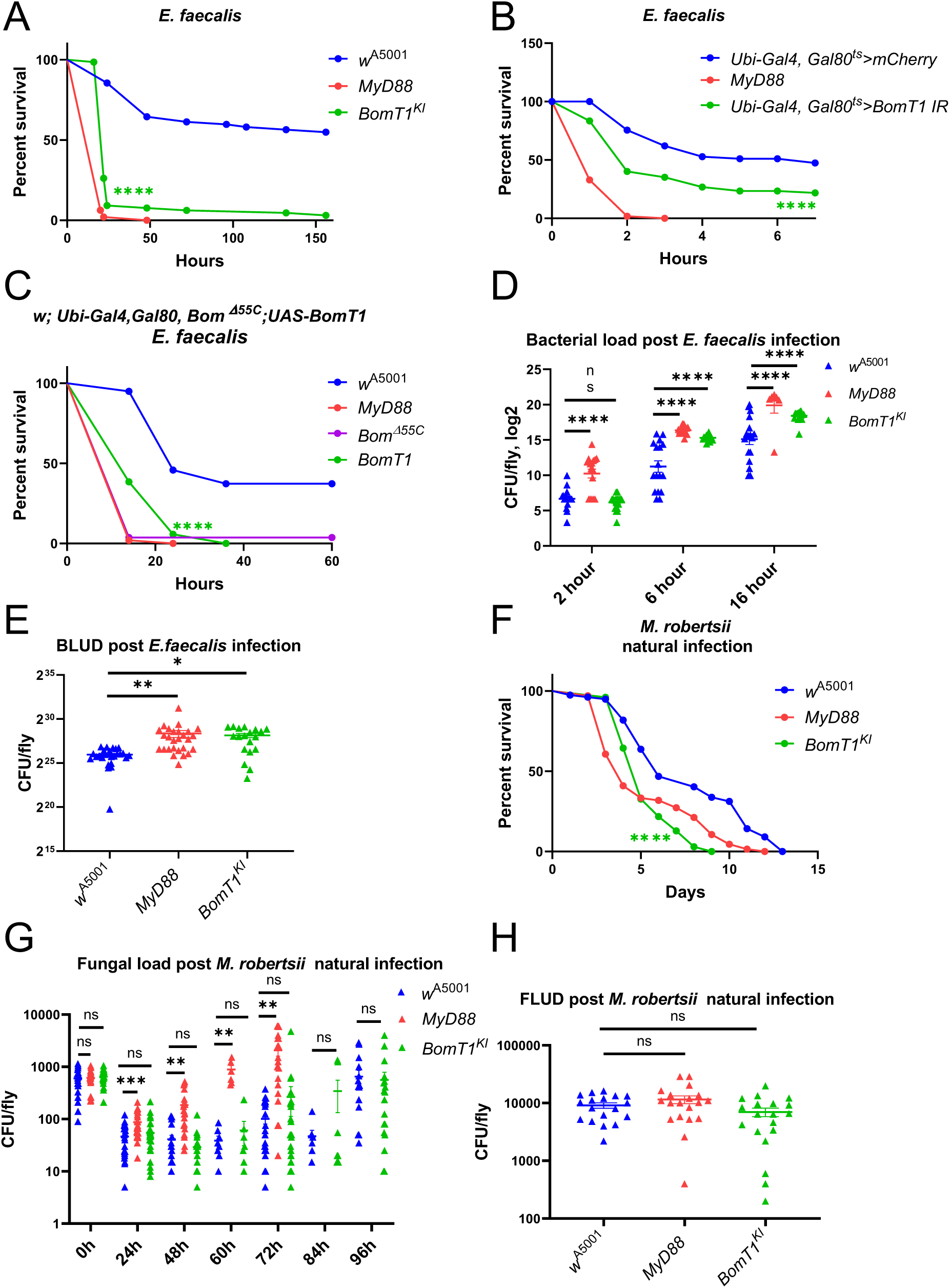
BomT1 partially mediates resistance to *E. faecalis* and resilience to *M. robertsii* infections. **(A-B)** Survival curves of *BomT1^KI^* mutant **(A)** or ubiquitous silencing of *BomT1* at the adult stage **(B)** flies after *E. faecalis* (OD_600_=0.1, 4.6 nL/fly) injection. **(C)** survival curves of flies ubiquitously overexpressing *BomT1* at the adult stage in *Bom^Δ55C^* background challenged by *E. faecalis* injection. **(D)** Bacterial load of *BomT1^KI^* mutant, *w* [A5001] and *MyD88* single flies post *E. faecalis* injection at 2h, 6h, and 16h. **(E)** Bacterial load upon death (BLUD) of *BomT1^KI^* mutant, *w* [A5001] and *MyD88* flies post *E. faecalis* injection. **(F)** Survival curves of *BomT1^KI^* mutant and *w* [A5001] flies after *M. robertsii* (10^5 spores/mL, 5mL/group) natural infection. **(G)** Fungal load of *BomT1^KI^* mutant, *w* [A5001] and *MyD88* flies post *M. robertsii* natural infection at different time points. **(H)** Fungal load upon death (FLUD) of *BomT1^KI^* single mutant flies, *w* [A5001] and *MyD88* flies post *M. robertsii* natural infection. Note: each experiment has been performed more than three times and pooled data are presented. **A, B, C, F**: Each experiment used biological triplicates of 20 flies. **D** and **G**: The data are presented as means ± SEM and analyzed using the ANOVA (one-way) with Dunnett’s multiple comparisons test; **E** and **H**: The data are presented as means ± SD and analyzed using the Unpaired t-test. * p<0.05; ** p<0.01; *** p<0.001; **** p<0.0001; ns, no significant difference.

We next investigated whether the *E. faecalis* burden was altered in *BomT1* KI single flies. As shown in Fig. 2D, the bacterial load was higher than in wild-type flies from six hours onwards, implying that *BomT1* is required for resistance. We also checked the bacterial load upon death (BLUD) of single flies and found that both *MyD88* and *BomT1* displayed an increased BLUD in the *w*[A5001] background (Fig. 2E), suggesting that more bacteria are needed to kill the *BomT1* and *MyD88* immuno-deficient flies. It is not clear however whether this finding reflects an increased resilience to *E. faecalis* infection.

### BomT1 may be involved in resilience to M. robertsii natural infection

We found that *BomT1* KI mutants were significantly more susceptible to *M. robertsii* natural infection, but not to the injection of its spores (Fig. 2F, Table 2). Of note, in contrast to the *E. faecalis* sensitivity phenotype, *BomT1*-silenced flies did not display any increased susceptibility to *M. robertsii* natural infection. This may reflect a hypomorphic effect of the RNAi approach, even though the silencing appeared rather strong at the transcript level (Fig. S7A). *M. robertsii* would also need to be more sensitive than *E. faecalis* to any remaining BomT1 in these silenced lines. In contrast to *MyD88* flies, the fungal load in *BomT1* single flies did not appear to vary at any time point (Fig. 2G). In terms of fungal load upon death (FLUD), there was no measured difference between wild-type and immunodeficient flies (Fig. 2H). Thus, our data are compatible with the hypothesis that BomT1 functions in resilience against *M. robertsii* natural infection.

**Table 2:**
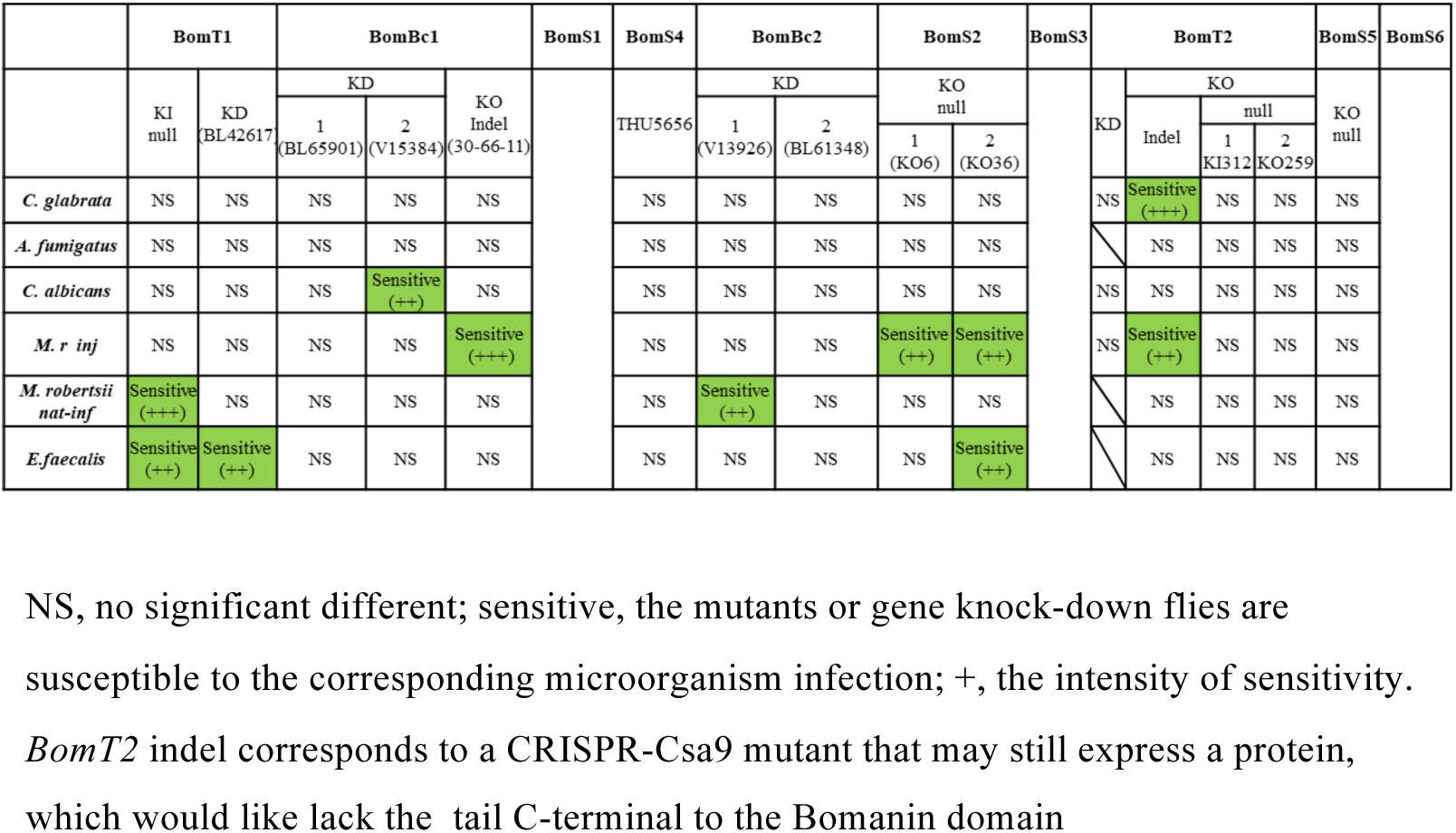
Loss-of-function *Bomanin* genes susceptibility to specific infections: a summary of *55C* locus *Bomanins*. Note: ns, not significant; sensitive, the mutants or single Bomanin knock-down flies are susceptible to the corresponding microorganism infection; ‘+’, the intensity of sensitivity. *BomT2* indel corresponds to a CRISPR-Cas9 mutant that may still express a protein, which would lack the tail C-terminal to the Bomanin domain.

Finally, *BomT1* overexpression in wild-type but not in *Bom^Δ55C^* flies provided a moderate degree of protection against the entomopathogenic fungus, suggesting it may act in concert with another *55C Bom*. In this respect, we discovered that the overexpression of *BomBc1*, *BomS4*, *BomS2* or *BomT2* protected flies from *M. robertsii* natural infection to the same extent than *BomT1* overexpression in wild-type flies (Table 1). Furthermore, *BomBc2*, *BomS3*, and especially *BomS1* overexpression provided an even stronger level of protection (Fig. EV3D-F, Table 1).

### BomS2 may be involved in resilience against injected M. robertsii

In opposition to *BomT1*, we found that the two *BomS2* null mutants were sensitive to injected *M. robertsii* spores but not to a natural infection with the same pathogen (Fig. 3A, Table 2). While we measured a somewhat increased fungal load at three days in *BomS2^ΔKO6^* mutant flies, we failed to reproduce this observation in *BomS2^ΔKO36^*flies as well as in *BomS2^ΔKO6^/ BomS2^ΔKO36^* trans-heterozygous flies (Fig. 3B-D). Thus, *BomS2* may be involved in resilience against this infectious challenge.

**Figure 3.**
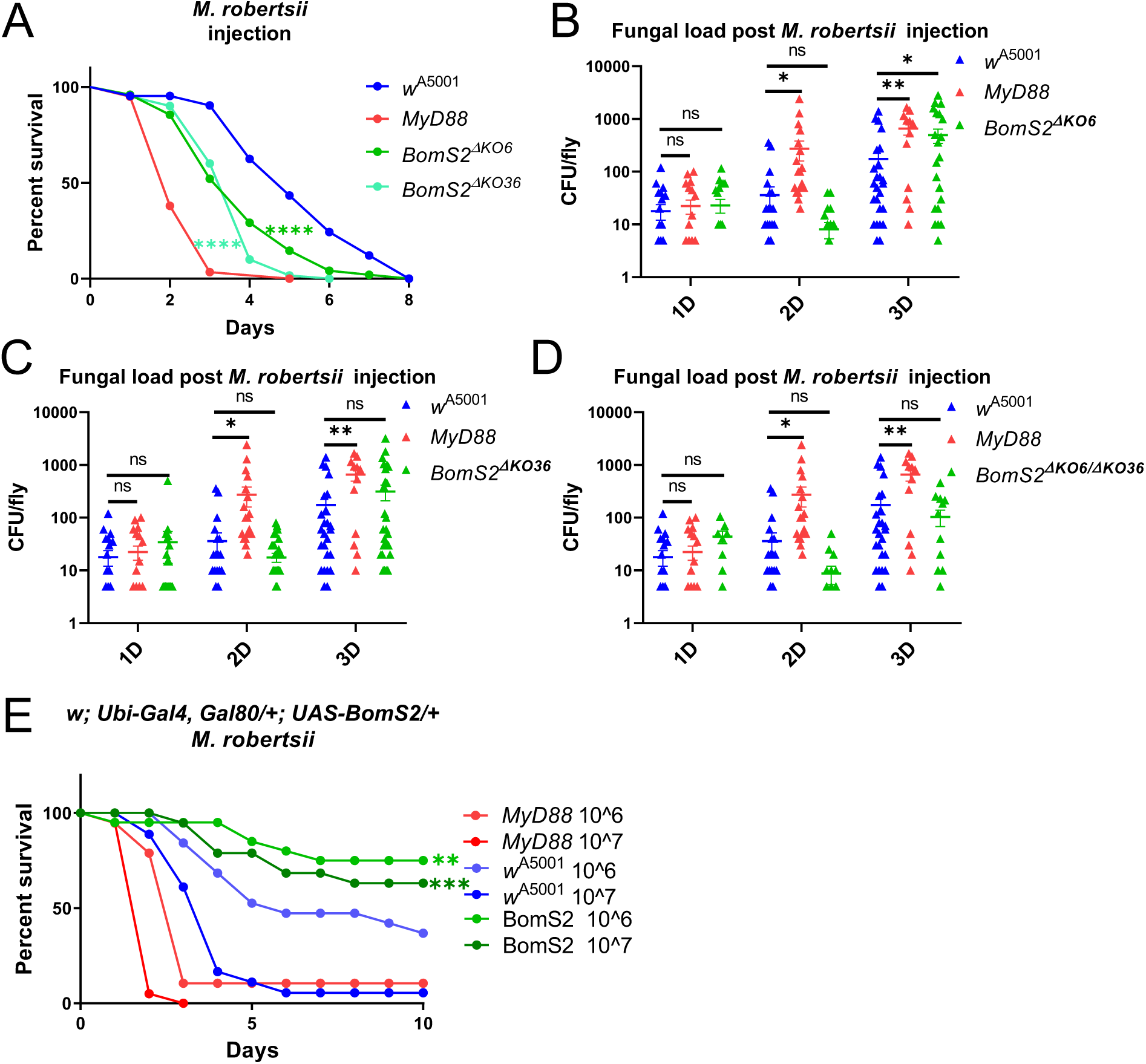
BomS2 also partially mediates resilience to injected *M. robertsii* spores. **(A)** Survival curves of two *BomS2* null mutants, *BomS2^ΔKO6^* and *BomS2^ΔKO36^* after *M. robertsii* (10^7 spores/mL, 4.6nL/fly) injection. **(B-D)** Fungal load of *BomS2^ΔKO^* **(B)**, *BomS2^ΔKO36^* **(C)**, and transheterozygous *BomS2^ΔKO6^/BomS2^ΔKO36^* mutant flies post *M. robertsii* injection at different time points **(D)**. **(E)** Survival experiments of *BomS2-*overexpressing flies in a wild-type background to *M. robertsii* injection. Note: each experiment has been performed more than three times and pooled data are displayed. **A** and **E**: Each experiment used biological triplicates of 20 flies. **B**, **C** and **D**: ANOVA, multiple comparisons, Dunnett’s multiple comparisons test. * p<0.05; ** p<0.01; *** p<0.001; **** p<0.0001.

The overexpression of *BomS2* in wild-type but not in *Bom^Δ55C^* flies protected them from the effects of *M. robertsii* injection at two inoculum doses (Fig. 3E, Table 1). Actually, we noted a similar level of protection conferred by the overexpression, also in adult wild-type flies, of *BomBc1* or of *BomS1* whereas that of *BomS4* or *BomS3* offered a more limited level of defense (Fig. EV4). Of note, *BomBc1* expresses distinct polymorphism isoforms in *w^1118^* and in *Canton-S* wild-type background (Fig. S2D). While the overexpression of the former enhanced the survival solely against injected *M. robertsii*, that of the latter did so only upon natural infection, an indication of the specificity of the roles of these two isoforms depending on the infection route of this pathogen (Table 1).

Unexpectedly, we detected that *BomS2^ΔKO36^* but not *BomS2^ΔKO6^* mutant flies displayed an increased sensitivity to *E. faecalis* (Fig. EV3A, Table 2), possibly in keeping with the report that *BomS2* silencing led to sensitivity to another, mild, Gram-positive bacterial pathogen, *Lysinibacillus fusiformis* (Smith *et al*., 2023).

### Bomanin-mediated host defenses against A. fumigatus, C. glabrata or C. albicans?

We have not identified any *Bom* gene with a clear-cut sensitivity phenotype to any of these three pathogens using the set of mutants/RNAi lines we have tested. The sensitivity of one *BomBc1* RNAi line to *C. albicans* was not confirmed in an independent line that appears as efficient in terms of silencing at the transcript level (Fig. S7B, S10B-C). Similarly, the susceptibility to a *C. glabrata* challenge of the *BomT2* indel line, which may still express a protein lacking the BomT2 tail, was not observed with the two null mutant lines (Fig. S10D-E).

We note that with respect to *C. glabrata* and *A. fumigatus* infections, the *Bom^Δ55C^*sensitivity phenotype can be rescued in general to a rather mild degree by the overexpression of a similar set of *Bom* genes, that is both *BomT* genes and *BomS* genes, with *BomS1* overexpression being able to protect the *Bom^Δ55C^* deficiency flies only against *A. fumigatus* whereas *BomS5* overexpression provided some defense to these flies only against *C. glabrata*.

Interestingly, the overexpression of *BomS1* in *MyD88* flies made them more sensitive to a *C. glabrata* challenge (Fig. S11A). It has been previously reported that Bombardier is involved in resilience by protecting the flies from the noxious effects of *BomS* genes when not secreted/stabilized in the hemolymph. Thus, BomS1 may contribute to the *Bbd* resilience phenotype. We have noticed in this respect that the continuous overexpression throughout the life cycle of the fly of *BomS1* led to a developmental phenotype characterized by Stubble-like bristles (Fig. S11B-D).

In contrast to the two infections above, the overexpression of only one *Bom* gene, the *w^1118^ BomBc1* isoform, protected to a mild degree *Bom^Δ55C^*flies against *C. albicans* infection (Fig. S10F).

### A role for Bomanins in preventing the dissemination of C. albicans throughout the fly body

Survival assays may not reflect the full palette of Bomanin functions. We reasoned that one aspect of infection is the dissemination of the pathogen away from its initial infection site. As a bright GFP-expressing strain of *C. albicans* was available, we examined how *C. albicans* behaved within the fly once injected in the thorax. As expected (DF, unpublished observations), the pathogenic yeast remained mostly at the injection site in *w* [A5001], yet they managed to form hyphae. In striking contrast, we observed a dissemination of *C. albicans* throughout the body of *Bom^Δ55C^* flies, with hyphae detected in all three tagmata (Fig. 4A).

**Figure 4.**
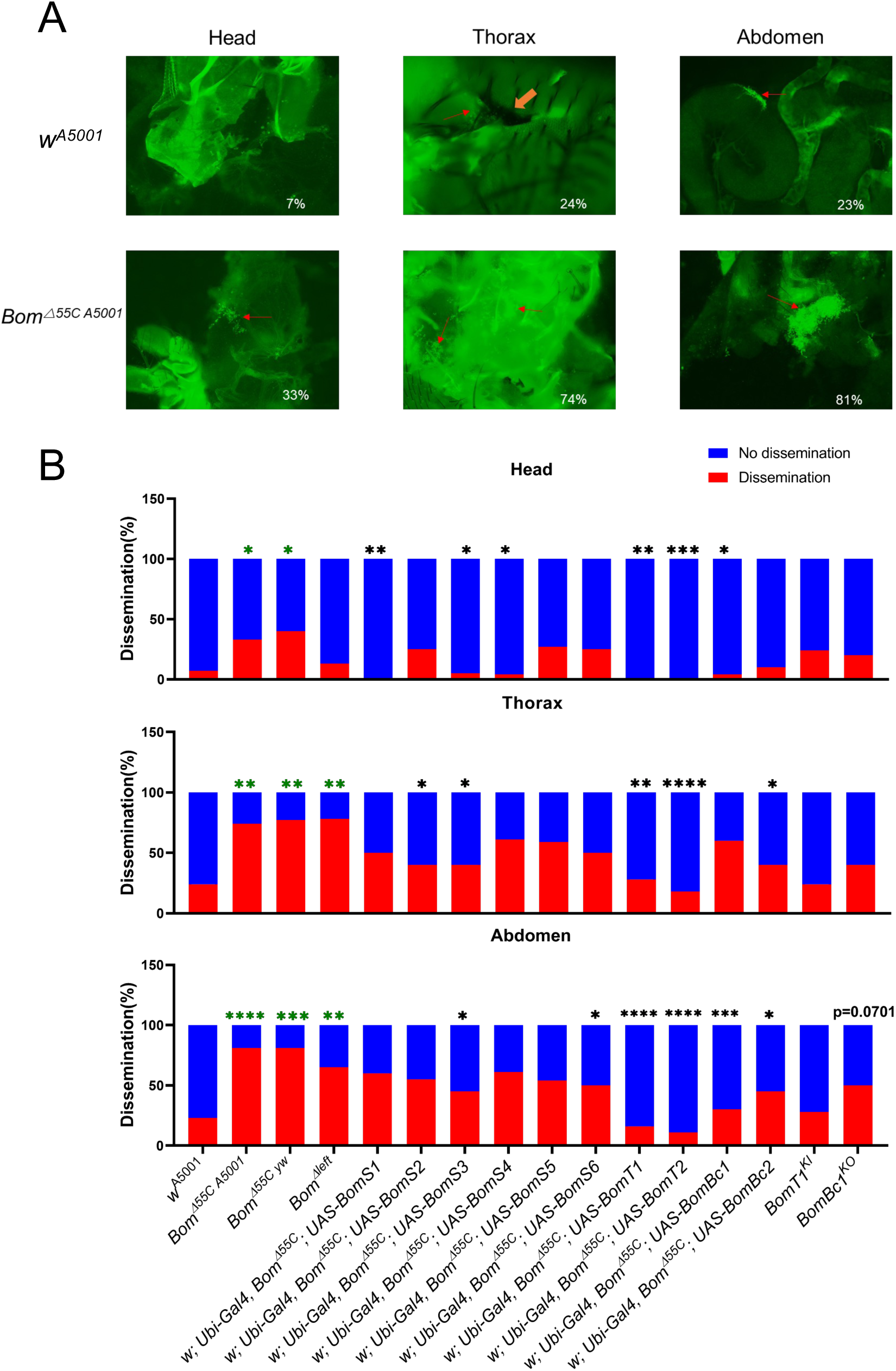
*55C Bomanins* prevent the dissemination of *C. albicans* throughout the fly body. **(A)** The conidia formation of infected flies with *C. albicans*. Red arrows: hyphae of *C. albicans*, yellow arrow: the melanized injection point. **(B)** Pooled data of *C. albicans* conidia formation in different flies at different fly sections. *w* [A5001]: wild-type flies. *MyD88*: *MyD88* mutant fly. *Bom^Δ55C^(A5001)*: *Bom^Δ55C^* backcrossed with *w* [A5001] fly. *Bom^Δ55C^*(yw): *Bom^Δ55C^* in initial *yw* background. *BomS1, BomS2, BomS3, BomS4, BomS5, BomS6, BomT1, BomT2, BomBc1, BomBc2*: overexpressed *BomS1, BomS2, BomS3, BomS4, BomS5, BomS6, BomT1, BomT2, BomBc1, BomBc2* flies individually in a *Bom^Δ55C^* background. *BomT1^ΔKI^*: *BomT1* mutant. *BomBc1^KO^*: *BomBc1* mutant, *BomBc1^indel^*. Note: each group contained at least 20 flies. **B**: The data are analyzed using Fisher’s exact test. * p<0.05; ** p<0.01; *** p<0.001; **** p<0.0001. Black *: comparison to *Bom^Δ55C^*; Green *: comparison to *w* [A5001].

We next complemented the *Bom^Δ55C^* phenotype by overexpressing each 55C *Bom* gene one-by-one. Both *BomT1* and *BomT2*, and to a lesser extent *BomBc1* prevented significantly the dissemination of *C. albicans* throughout the body of *Bom^Δ55C^* flies. Conversely, there was a trend for the *BomBc1* indel mutant but not for the *BomT1* KI mutant flies to allow some dissemination of *C. albicans* in an otherwise wild-type background (Fig. 4B).

### Implication of the 55C Bomanin cluster in the host defense against A. fumigatus mycotoxins

We have previously reported that *Bom^Δ55C^* are sensitive to the injection of two mycotoxins secreted by *A. fumigatus*, namely the ribotoxin protein restrictocin and the family of fumitremorgin/verruculogen compounds (Fig. S12). We had also reported that *A. fumigatus* mutants lacking the locus encoding restrictocin or lacking the fumitremorgin biosynthetic cluster were only mildly less virulent than the wild-type fungus in *MyD88*-immunodeficient flies. We have extended these observations to the *Bom^Δ55C^* and *Bom^Δleft^* fly lines. Whereas we still observed a reduced virulence in the *Bom^Δ55C^*line, as for *MyD88* flies, we no longer measured a significant difference in the virulence of the mycotoxin-defective fungi as compared to wild-type fungi in the *Bom^Δleft^*line (Fig. EV5A-D). This suggests that the four genes remaining in *Bom^Δleft^* flies, namely *BomS3*, *BomT2*, *BomS5*, and *BomS6* are sufficient to protect the flies against restrictocin and verruculogen/fumitremorgins.

We also tested directly the difference in sensitivity between *Bom^Δ55C^* and *Bom^Δleft^* after exposure to restrictocin, fumitremorgin B or verruculogen. Whereas verruculogen killed equally well both deletion lines, restrictocin and fumitremorgin B were not as toxic to the *Bom^Δleft^* as to the *Bom^Δ55C^* line (Fig. 5A-C), suggesting that the four remaining genes in *Bom^Δleft^* are not sufficient to confer protection against restrictocin and fumitremorgin B and that at least some of the six genes removed by this deficiency are required for host defense against these mycotoxins. We next injected restrictocin or verruculogen into *BomT1* KI mutant and the *BomBc1* indel mutant. We found that both mutants are as sensitive to restrictocin as *MyD88* mutants, whereas none of them displayed any enhanced susceptibility to a verruculogen challenge (Fig. 5D-G).

**Figure 5.**
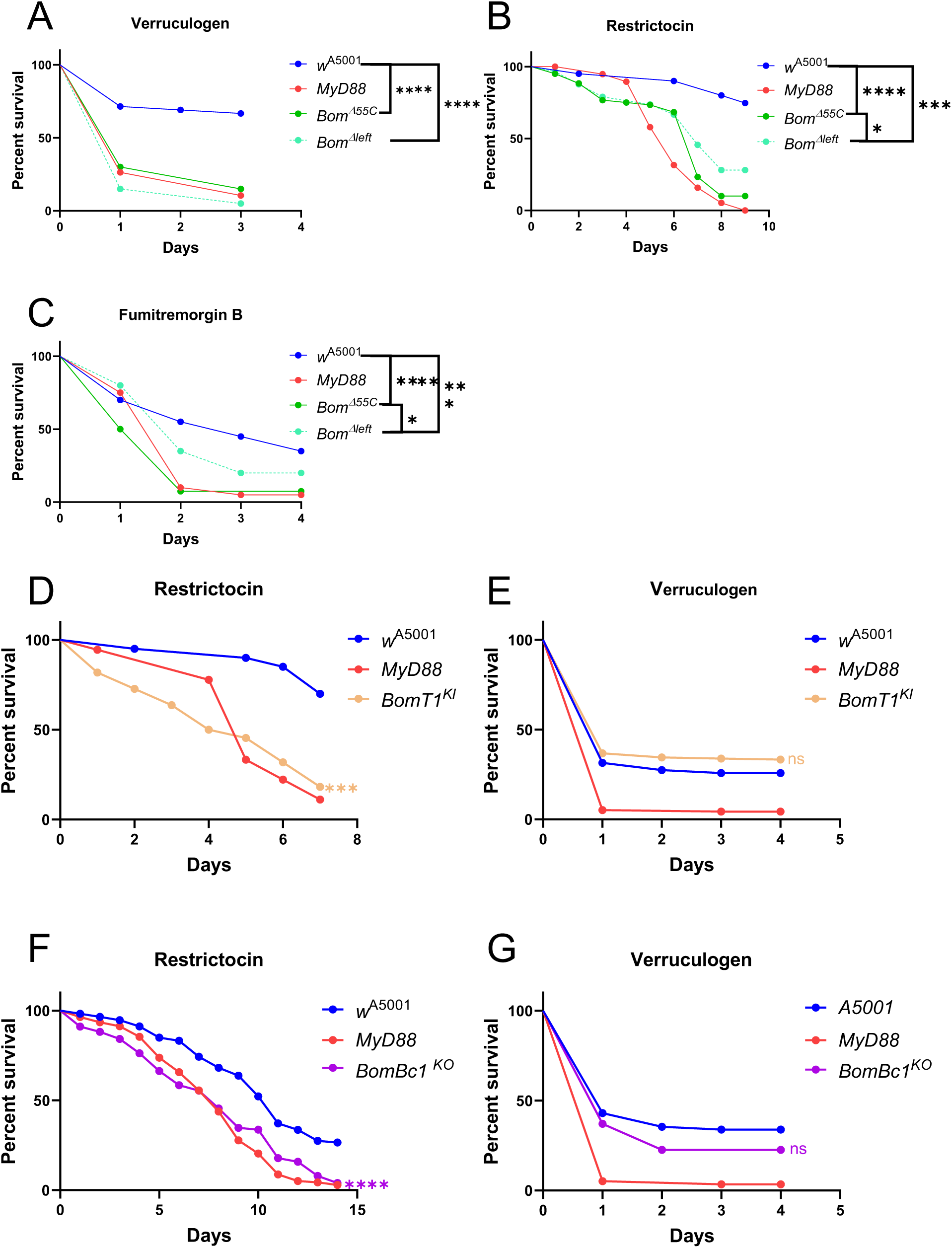
*55C Bomanins* confer protection against *A. fumigatus* mycotoxins. **(A-C)** Survival curves of *Bom^Δ55C^*, *Bom^Δleft^, MyD88,* and *w* [A5001] flies after commercially-available mycotoxins, Verruculogen (5mg/ml, 4.6 nL/fly) (A), Restrictocin (5mg/ml, 2 nL/fly) (B), and Fumitremorgin B (5mg/ml, 2 nL/fly) (C) injection. **(D-E)** Survival curves of *BomT1^KI^* flies after Restrictocin (D) and Verruculogen (E) injection. **(F-G)** Survival curves of *BomBc1^KO^* flies after Restrictocin (F) and verruculogen (G) injection. Note: **(A-G)** Each experiment has been performed more than three times and pooled data are presented. Except for **(E)** and **(G)**, each experiment used biological triplicates of 20 flies.

Finally, as a proxy to analyze the function of *BomSs* in the resilience to *A. fumigatus* mycotoxins, we tested *Bbd* mutants in which no BomSs are detectable in the hemolymph. We found *Bbd* mutant flies to be as sensitive to *A. fumigatus* as *MyD88* flies (Fig. S13A). They were somewhat sensitive to restrictocin, highly sensitive to verruculogen and not susceptible to a gliotoxin challenge (Fig. S13B-D). Of note, we formally cannot a contribution of *55C* BomTs to these phenotypes since it is unknown whether Bbd is required for the secretion/stability of BomTs in the hemolymph. We have previously documented the potential for *BomS3* (restrictocin only) and especially *BomS6* to rescue the sensitivity of *Bom^Δ55C^* flies to injected restrictocin or verruculogen (Xu *et al*., 2023).

## Discussion

In this article, we report a genetic dissection of the *55C Bomanin* locus using both loss-of-function and overexpression strategies. A major outcome of this study is the identification of BomT1 as being a major effector of host resistance against *E. faecalis* infection, albeit it likely functions together with other gene product(s) of the locus. Interestingly, the same gene appeared to be required in the host defense against *M. robertsii* natural infection; we failed to obtain evidence supporting a role for it in resistance against this infection. Likewise, we found that *BomS2* but not *BomS5* null mutants are required for protection against injected *M. robertsii* spores even though the fungal burden remained unaltered as compared to wild-type flies. We also report that two Bomanins are required for protection against a ribotoxin secreted by *A. fumigatus*, including *BomT1*, implying that this Bom peptide is involved both in resistance and resilience to infections. Finally, we have discovered a disconnect between fungal dissemination and fungal lethality, with again BomTs appearing to prevent to some extent the dissemination of *C. albicans* hyphae. Thus, the functions of *55C* Bomanins are complex and intertwined and involved in both the resistance and resilience facets of host defense.

### Limitations of the study

Even though we have generated five null mutants at the *55C* locus, we have failed to obtain any reagent affecting the expression of *BomS1, BomS3*, and *BomS6*; we obtained data only by silencing gene expression for *BomS4* and *BomBc2*. As regards *BomS4*, we have analyzed only one RNAi line, which however appears to be rather effective at silencing *BomS4* transcripts. Yet, we cannot exclude a hypomorphic effect or a potential accumulation of the basally expressed BomS4 peptide prior to the silencing of the transcripts at the adult stage. Thus, because we did not find any *BomS4* survival phenotype with the pathogens tested, off-target effects are not an issue; yet, a null mutant would be required to confirm this apparent absence of roles in the host defense against these bacterial and fungal pathogens.

The deletion of one gene in a locus may affect the neighboring one, as appears to be the case for *BomS5* for which *BomT2* expression appears to be severely affected. As regards *BomT1*, we have noted a reduced induction of *BomS2* expression, which is located 5 kB away. It is unlikely that a second-site mutation in the KI mutant would also occur upon silencing *BomT1*. We also note that *BomT1* null mutants affect the survival to *M. robertsii* natural infection but not to that of injected spores whereas the *BomS2* null phenotype is the converse.

Of interest, this study confirms that distinct host defenses are relevant according to the route of infection of *M. robertsii* (Wang, 2020). Notwithstanding, another limitation is that we have tested here only a limited panel of bacterial, yeast and filamentous fungal pathogens.

Finally, overexpression experiments have to be interpreted with caution as the degree of overexpression of the peptides reached may not be physiologically relevant; results from this approach should be taken as indications of the potential abilities of the overexpressed gene. Nevertheless, we report in this study some specificity in the effects of overexpressed Bomanins, with some overlap in survival phenotypes to infections between some but not all short Bomanins, in contrast to a previous study on the redundant functions of BomS against *C. glabrata* (Lindsay *et al*., 2018).

### Role of BomT1 in the resistance to E. faecalis infection

Clemmons *et al*., had reported that the *55C* locus is required for resistance against *E. faecalis* (Clemmons *et al*., 2015). We found that *BomT1* mutants displayed a susceptibility to this bacterial pathogen that was almost as pronounced as that displayed by *MyD88*, suggesting that it is a major mediator of resistance to *E. faecalis* since the bacterial burden is increased in this mutant. Yet, it may not act alone against this bacterium since we observed only a moderate rescue of the *Bom^Δ55C^* sensitivity phenotype by its overexpression. We note that *Bom^Δleft^* are also as susceptible to this infectious challenge, suggesting that the four-rightmost genes of the locus are not sufficient to mediate protection on their own. Interestingly, *BomS1* overexpression also somewhat rescues *Bom^Δ55C^* flies and it would be interesting to test whether their joint overexpression in this immunodeficient context would bring the protection to wild-type levels. Unexpectedly, we found that *BomBc2* provided a degree of protection against *E. faecalis* when overexpressed in wild-type but not *Bom^Δ55C^* flies, possibly reflecting a positive interaction with another *55C Bomanin* gene product, possibly *BomT1*. A *BomBc2* null mutant will be required to clarify the function of this gene in host defense against infections. A weak phenotype of susceptibility to *E. faecalis* has previously been reported upon silencing *BomBc1* but not *BomT1*, *BomS1*, and *BomS4* (Chapman *et al*., 2020), in contrast to the results presented in our study (Table 2). Our analysis indicates that specific Bomanins play a role in host defense against *E. faecalis* and thus are not compatible with the hypothesis that it is the overall amount of Bomanin peptides that is the relevant parameter for controlling microbial infections (Clemmons *et al*., 2015; Lindsay *et al*., 2018).

### Involvement of 55C genes in resilience to infections

*BomS2* behaves much as *BomT1* mutants with no altered fungal burden after a challenge with *M. robertsii* spores thus arguing against a role in resistance. It is an open possibility that *BomT1* and *BomS2* rather function in resilience to secreted virulence factors. We have reported previously that some BaraA-derived peptides provide a degree of protection against Destruxin A, a toxic hexadepsipeptide secreted by the fungus (Huang *et al*., 2023). It remains unclear whether the *55C* Bomanin locus is involved in resilience to Destruxin A as we have obtained contrasted results in our experiments. However, the number of secreted proteins, many of them proteases, and of secondary metabolites is so high that the two Bomanins may protect against other virulence factors (Gao *et al*, 2011). Interestingly, some virulence factors able to inhibit the activation of the Toll pathway by either the GNBP3 sensor or the PSH protease bait have recently been identified (Lu *et al*, 2024; Tang *et al*, 2025). However, as pointed earlier (Clemmons *et al*., 2015), the basal expression levels of many *Bomanins* in adult fat body or carcass is high, especially those of *BomS1, BomBc2*, *BomS2*, *BomS3*, *BomT2*, and *BomS6*, which might be sufficient to provide a degree of protection despite the pathogen attack on the activation of the Toll pathway.

Nevertheless, the strongest evidence we have obtained for a role of Bomanins in resilience against a mycotoxin is the sensitivity of both *BomT1* and *BomBc1* to a challenge by the *A. fumigatus* mycotoxin restrictocin (this work).

Of note, a recent study has documented a role for *BomS2* in the host defense against *Lysinibacillus fusiformis*, a mild Gram-positive pathogen. With respect to *E. faecalis*, we did observe a mutant phenotype for only one of two null mutants (Fig. EV3A, Table 2).

### The dissemination of C. albicans within the Drosophila body may not be essential for its pathogenicity

*C. albicans* is a dimorphic yeast able to form hyphae. The transition between the yeast to filamentous forms has been reported to be important for its virulence and it is believed that hyphae allow it to invade tissues thereby promoting its pathogenicity. Here, we find that *Bom^Δ55C^* mutants are highly permissive to the dissemination of the fungus, in keeping with its sensitivity to this infection. The rescue experiments of *Bom^Δ55C^* by single *Bom* gene overexpression revealed that *BomT1* and *BomT2* strongly prevent the dissemination of *C. albicans* within the fly yet do not allow it to better survive the infection. Since *BomT1* or *BomT2* overexpression did not protect *Bom^Δ55C^* mutants from *C. albicans* infection (Table 1), these findings suggest that *C. albicans* can kill its host in the absence of a widespread dissemination.

Thus, this pathogen may kill *Bom^Δ55C^* hosts through secreted virulence factors, a situation akin to that recently described for *A. fumigatus* infection (Xu *et al*., 2023). Yet, protection against these secreted virulence factors would necessarily be mediated by multiple *55C* Bomanins since the overexpression of single *55C* Bomanin genes did not protect *Bom^Δ55C^* flies from *C. albicans* infection (Table 1).

In contrast, the overexpression of a specific *BomBc1* isoform also inhibited the dissemination of *C. albicans* and conversely the deletion of the *BomBc1* locus nearly led to a significant dissemination of the fungus (Fig. 4B). Of note, we cannot totally exclude that we would have been able to detect with a high sensitivity any dissemination of *C. albicans* under the yeast form. Nevertheless, our data suggest that some long Bomanin peptides may interfere with filamentation. Interestingly, it has been reported that Daisho peptides, which are related to Bomanins (Clemmons *et al*., 2015), have been shown to bind to hyphae of the mold *Fusarium oxysporum* against which they are active (Cohen *et al*., 2020).

This study reveals that distinct Bomanins have different activities in host defense against infections, ranging from antimicrobial activities against a prokaryote pathogen, *E. faecalis*, against eukaryotic pathogens including yeast and filamentous fungi and also provide protection against secreted virulence factors. Strikingly, BomS6 is able to protect flies from the action of a protein toxin, Restrictocin, and a secondary metabolite, verruculogen that targets a maxi-potassium channel by binding to a site within its transmembrane domain. At first sight, it is difficult to envision how this family of related peptides manages to function in such diverse facets of host defense. We would like to speculate here that there is a potential common target of this family of Toll pathway effectors, namely cytoplasmic membranes. Like several AMPs such as Cecropins, antimicrobial Bomanins such as BomT1 may interact with the bacterial membrane and possibly directly lead to its lysis, likely in association with other factors such as other Bomanins. A similar process may be at play as regards the fungicidal activity of BomS or action on hyphae of BomTs/BomBc1. With regard to the protection against mycotoxin, we note that a remarkable feature of Restrictocin is its ability to cross membranes. Future studies will tell whether BomS6 is able to modify the host cell permeability to the ribotoxin and whether the modification of the biophysical properties of neural membranes induces a conformational change in the Slowpoke receptor that would mask its verruculogen binding site, thereby allowing the host to recover from tremors induced by this neuromycotoxin (Xu *et al*., 2023).

## Material and methods

### Fly stocks and maintenance

#### Fly incubation

Flies were maintained under controlled environmental conditions (25°C, 65% RH) in standard incubators. The nutritional medium was prepared by combining the following components per production batch: 1.2 kg cornmeal (Priméal), 1.2 kg glucose (Tereos Syral), 1.5 kg yeast (Bio Springer), 90 g nipagin (VWR Chemicals) diluted into 350 mL ethanol (Sigma-Aldrich), 120 g agar-agar (Sobigel). Ultrapure water was added to obtain a 25 L batch of food.

#### Indel mutants and RNAi lines

The following mutant lines were used: the *Bom^Δ55C^*and *Bom^Δleft^* lines were a kind gift of Prof. Steven Wasserman. *BomT2^Δtail^* and *BomBc1^indel^* were generously provided by Prof. Jiyong Liu. The *BomT1^KI^* line was contructed by Wellgenetics (Taipei, Taiwan). Other KO mutants were constructed by ourselves using CRISPR-Cas9 technology (see Supplementary Material). All mutant strains were isogenized in a *w*[A5001] background (Thibault *et al*., 2004). The following knockdown strains obtained from the Bloomington, Vienna, and Tsinghua stock centers were utilized: *BomT1* (BL42617); *BomBc1* (BL65901); *BomBc2* (VDRC13926) and BomBc2 (BL61348); *BomT2* (VDRC103059); *BomS4* (THU5656). All the RNAi lines were crossed to a *w; pUbi-Gal4, pTub-Gal80*^ts^ (BDSC30140) driver lines at 18°C; hatched adults were placed at 29°C for 5 days to induce RNAi expression. **Transgenic lines**. The transgenic lines expressing single *Bom* genes of the *55C* locus under the *pUAS-hsp70* promoter control were generated (see Supplementary Material) and checked by sequencing (Xu *et al*., 2023). The transgenic flies were crossed to a *w; pUbi-Gal4, pTub-Gal80^ts^*driver line, in a homozygous *MyD88*, *Bom^Δ55C^* mutant or *w*^A5001^ background. The expression of the transgenes was checked by RT–qPCR and mass spectrometry analysis on collected hemolymph of single flies.

### Quantitative RT-PCR

Total RNA was extracted from the whole-body of adult flies using TRIzol® reagent (Invitrogen), following the manufacturer’s protocol. For each biological replicate, five age- and sex-matched flies were pooled to minimize individual variability. RNA integrity was confirmed via NanoDrop. Complementary DNA (cDNA) was synthesized from 1 μg of total RNA using the iScript™ cDNA Synthesis Kit (Bio-Rad), with random hexamer primers, according to the manufacturer’s instructions. The primers used in quantitative PCR were listed in Table S5.

### Mass spectrometry

Single fly hemolymph was collected from individual flies and immediately transferred to 2 µL of 0.1% trifluoroacetic acid (TFA) solution on ice to prevent protein degradation and maintain sample integrity. The 4-hydroxy-α-cyanocinnamic acid (4-HCCA) sandwich spotting matrix method was employed for sample loading. Two distinct matrix solutions were prepared: Matrix 1: A saturated solution of 4-HCCA (20 mg/mL) in 100% acetone. Matrix 2: A solution containing 20 mg/mL 4-HCCA in a 2:1 acetonitrile/0.1% TFA mixture. For each hemolymph sample from a single fly, 0.6 µL of the sample was loaded onto the MALDI target plate. The sample was directly applied to a dried bed of 0.5 µL of Matrix 1 and the sample was overlaid with 0.4 µL of Matrix 2 to ensure optimal crystallization and ionization efficiency during MS analysis. After air drying under ambient condition, the prepared samples were analyzed using a MALDI-TOF mass spectrometer. The instrument was calibrated with standard peptide/protein mixtures to ensure accurate mass measurements. Data acquisition was performed in positive ion mode over a mass range of 800–20,000 Da, with a laser intensity optimized for the detection of small peptides and proteins present in the hemolymph.

### Microbial culture and infection

*Enterococcus faecalis*. *E. faecalis* strains (*ATCC 19433*) were maintained on Luria-Bertani agar (LBA) plates stored at 4°C for at most a month. For experimental replicates, single colonies were isolated under sterile conditions and inoculated into 5 mL LB broth. Cultures were incubated at 37°C with shaking (220 rpm) for 6-8 hours to reach mid-logarithmic phase (OD_600_ = 0.6-0.8, measured via Nanodrop spectrophotometer). Cells were collected by centrifugation (7,500 rpm, 2 min, 4°C) and resuspended in phosphate-buffered saline (PBS). This washing procedure was repeated three times to remove residual culture media. Final bacterial suspensions were adjusted to OD_600_=0.1 concentration, ensuring experimental consistency across biological replicates. 4.6 nL of standardized bacterial suspension was delivered per fly via thoracic microinjection

*Candida albicans* (*CAM15.4*) and *C. glabrata* (*ATCC2001*) were maintained on yeast extract peptone dextrose (YPD) after two rounds on agar plates at 29°C. Single colonies were taken using a sterile tungsten needle for infections, 3-5 days old adult flies were anesthetized and pricked in the thorax region with the needle containing fungal cells (colony pricking assay). Mock-infected controls were performed using needles dipped into sterile PBS.

*Metarhizium robertsii* (*2575*) was propagated on Potato Dextrose Agar (PDA; BD Difco™) plates at 25°C for 7-14 days to ensure optimal sporulation. Spores were collected by gently scraping plate surfaces with PBST (PBS + 0.01% Tween-20) for injection experiments or sterile ddH_2_O + 0.01% Tween-20 for natural infection assays. Suspensions were filtered through Miracloth to remove mycelial debris and quantified using a hemocytometer. For injection experiments, 4.6 nL of standardized conidial suspension (10^7 spores/mL) was delivered per fly via thoracic microinjection. For natural infection, 5 mL of diluted conidial suspension (10^5 spores/mL) was applied to groups of 20 flies.

*Aspergillus fumigatus*. *A. fumigatus* strain maintenance and infection procedures were performed as previously described (Xu *et al*., 2023).

*Micrococcus luteus. M. luteus* was cultivated in Luria-Bertani Broth at 37°C under aerobic conditions for 24 hours. Bacterial cells were pelleted by centrifugation at 3,000 rpm for 10 minutes, followed by two successive washing cycles in phosphate-buffered saline (PBS). The pellet was resuspended in 1 mL PBS after each centrifugation step to ensure complete removal of culture medium components. Needles were dipped in a concentrated pellet prior to pricking flies.

### Bacterial/ fungal load in single flies

Individual infected flies were processed for microbial enumeration using a modified protocol as follows: single flies were transferred to 1.5 mL microcentrifuge tubes containing 20 µL phosphate-buffered saline with Tween-20 (PBST) and two 3 mm stainless steel beads. Samples were homogenized by mechanical disruption using a Mixer Mill (Retsch MM 300) at 30 Hz for 2 min. Homogenates were subjected to serial dilutions (10⁻¹ to 10⁻⁵) in PBST based on pre-determined time-point-specific infection kinetics. Aliquots (50 µL) of each dilution were spread-plated in duplicate onto potato dextrose agar (PDA) plates for fungi or LBA ones for bacteria, with one fly equivalent per plate. Plates were incubated at organism-specific temperatures (29°C for fungi, 37°C for bacteria) in a humidified incubator. Fungal colonies were enumerated after 48 h incubation, while bacterial colonies were counted after 24 h. The limit of detection was established at 10 CFU/fly based on the lowest dilution yielding ≥30 colonies per plate. Flies died in 30 minutes after injection were collected for Bacteria load upon death (BLUD) or fungi load upon death (FLUD) analysis.

### Mycotoxins preparation and injection

Mycotoxin standard solutions were prepared as described previously (Xu *et al*., 2023). Commercially available mycotoxin powders (Restrictocin in PBS, verruculogen in dimethyl sulfoxide (DMSO), and fumitremorgin B in DMSO) were dissolved to generate stock solutions. Working concentrations were prepared by serial dilution in PBS or DMSO. Stock solutions were stored at −20°C in amber vials protected from light, while fresh working solutions were prepared daily.

### Survival tests

5-7 day-old adults were utilized to ensure uniform physiological maturity. Flies were housed in vials containing standard cornmeal-agar medium at a density of 20 individuals per vial, with three biological replicates maintained for each experimental condition. During infection studies, daily survival rates were monitored by transferring surviving flies to fresh vials and recording mortality.

### *In vivo* phagocytosis

Phagocytosis of *Bom^Δ55C^*mutant flies was measured following our previously published protocol (Liégeois et al, 2020).

## Statistical analysis and reproducibility

All survival data were analyzed using GraphPad Prism 9.0 software (GraphPad Software, San Diego, CA, USA). Survival curves were compared using the log-rank (Mantel-Cox) test unless otherwise specified in the figure legends. For quantitative analyses of microbial burden, gene expression levels, and GFP fluorescence intensity, unpaired, non-parametric Mann-Whitney statistical tests or Student t-test were employed as appropriate; dissemination data were anlyzed using Fisher’s exact test. Significance values: *P < 0.05; **P < 0.01; ***P < 0.0003; ****P < 0.0001; ns, not significant.

A list of reagents is available as Supplementary Table 1.

## Data availability

This study includes no data deposited in external repositories.

## Acknowledgements

We thank Anne Beauvais and Jean-Paul Latge for the *A. fumigatus* strain used in this study, Bruno Lemaitre, Jiyong Liu, Steven Wasserman, and the Guangzhou Drosophila Resource Center for fly stocks. Stocks obtained from the Bloomington Drosophila Stock Center, Vienna, and *Tsinghua* stock centers were also used in this study. We gratefully acknowledge the contributions of Miriam Yamba for expert technical help. YL was respectively partially funded through the Sino-Foreign cooperative graduate education project of Guangzhou Medical University, the International Training Plan for young outstanding scientific research talents of Guangdong Province, and Overseas Postdoctoral Talent Support Program of Guangdong Province. This work was supported by the Association Platform BioPark of Archamps on its Research & Development budget (PB), from the National Natural Science Foundation of China Project (#32370931), the China High-end Foreign Talent Program, and the 111 Project (#D18010; China) to DF.

## Author contributions

**Yanyan Lou**: Conceptualization; resources; formal analysis; validation; investigation; methodology; writing – original draft; writing – review and editing. **Bo Zhang**: Conceptualization; resources; formal analysis; investigation; methodology; writing – original draft; writing – review and editing. **Zhiyuan Zhang, Yingyi Pan, Jianwen Yang, Lu Li, Jianqiong Huang**: Resources; investigation; writing – review and editing. **Zihang Yuan**: investigation; writing – original draft; writing – review and editing. **Samuel Liégeois**: resources; formal analysis; validation; investigation; methodology. **Philippe Bulet**: Resources; formal analysis; validation; investigation; methodology; performed and analyzed the mass spectrometry analysis; **Rui Xu**: funding acquisition; designed, analyzed the mycotoxins experiments. **Zi Li**: Conceptualization; resources; funding acquisition. **Dominique Ferrandon**: Conceptualization; resources; supervision; funding acquisition; validation; methodology; writing – original draft; project administration; writing – review and editing.

## Expanded view figure legends

**Figure EV1:**
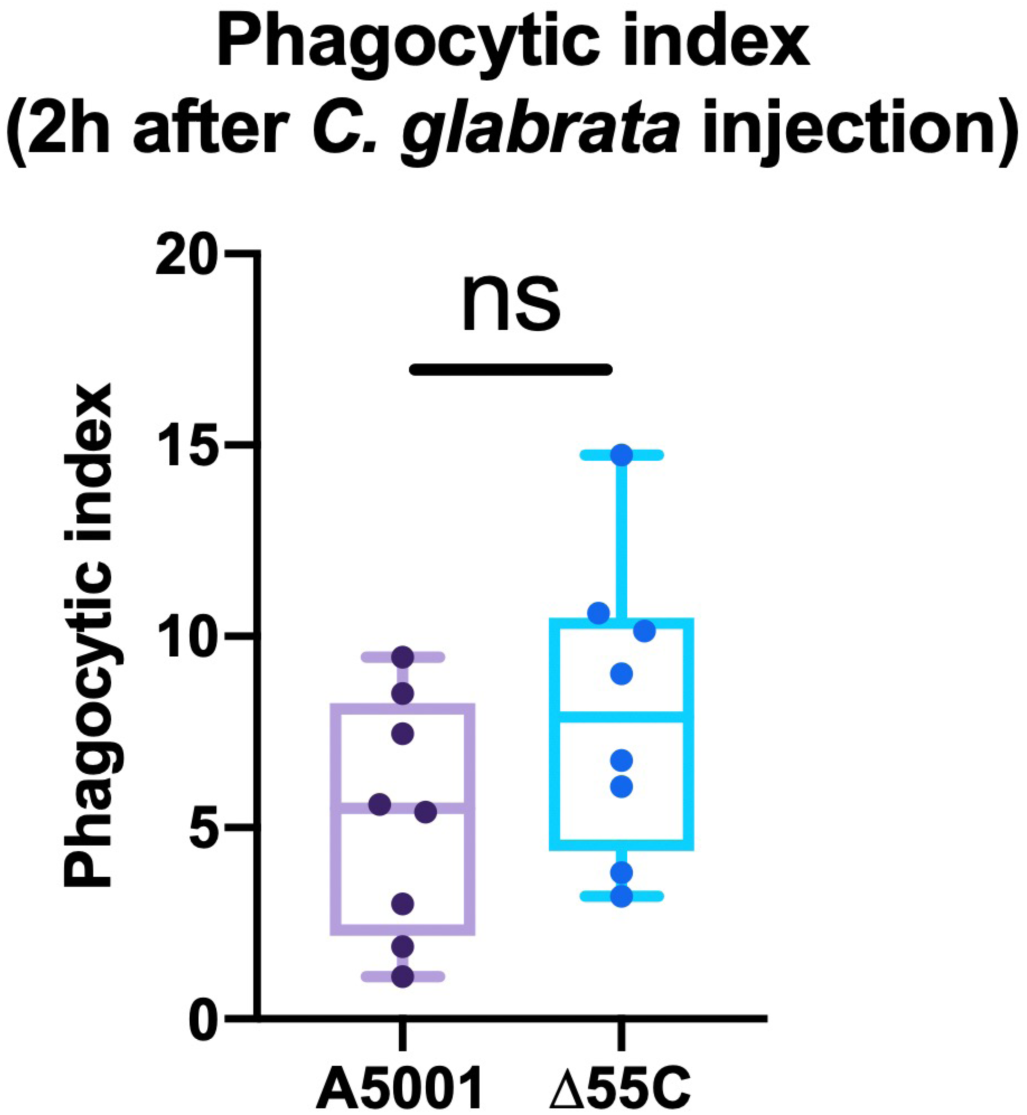
*Bomanins* at *55C* locus are not involved in phagocytosis. Complementary to Fig. 1 6,000 live yeast cells of *C. glabrata* were injected into *w* [A5001] wild-type flies and *Bom^Δ55C^*flies (Δ55C) and their phagocytic index was monitored 2 hours after injection. Mann-Whitney test.

**Figure EV2:**
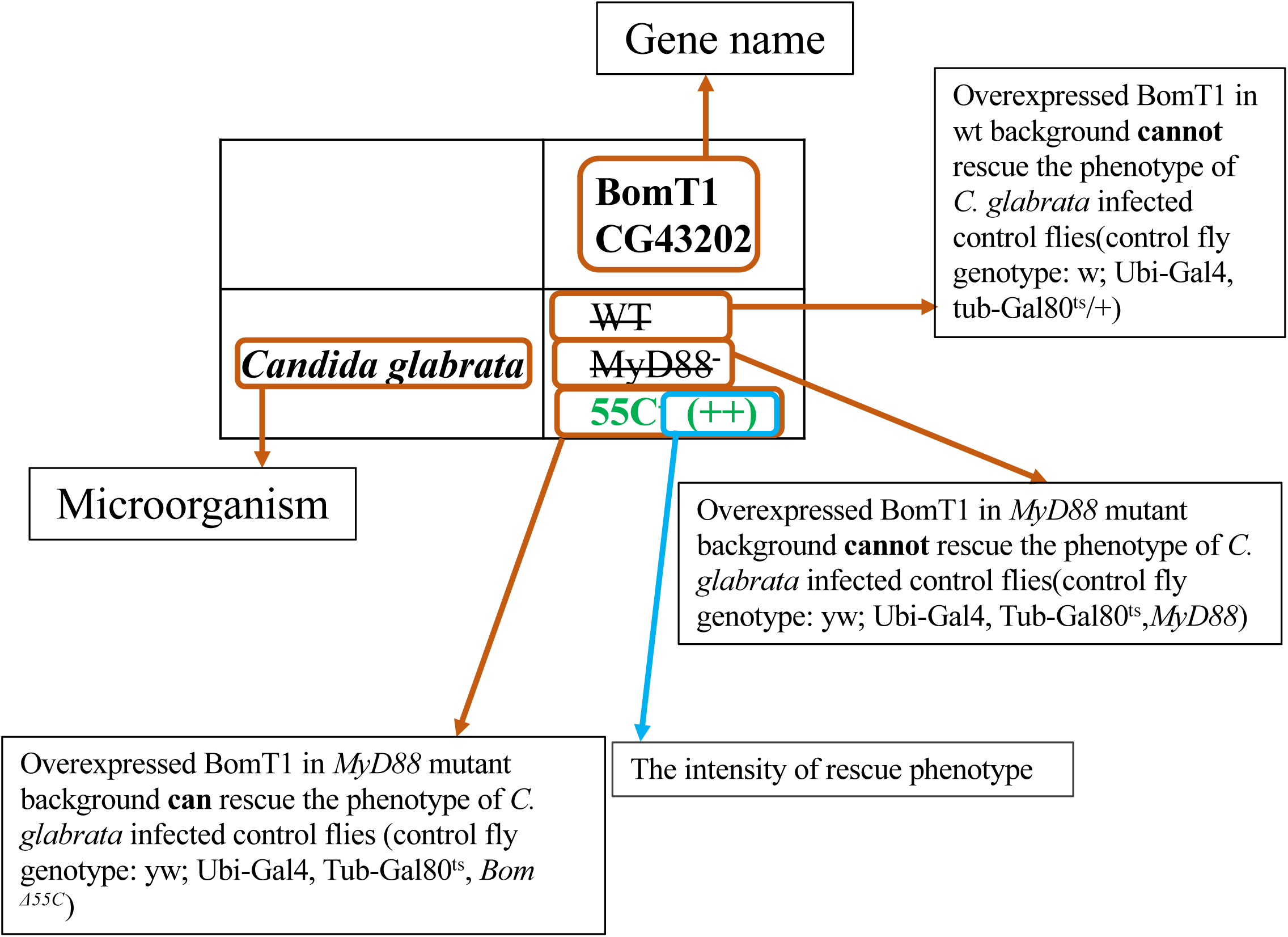
A key annotation of every single cell for Table 1. Complementary to Table 1.

**Figure EV3:**
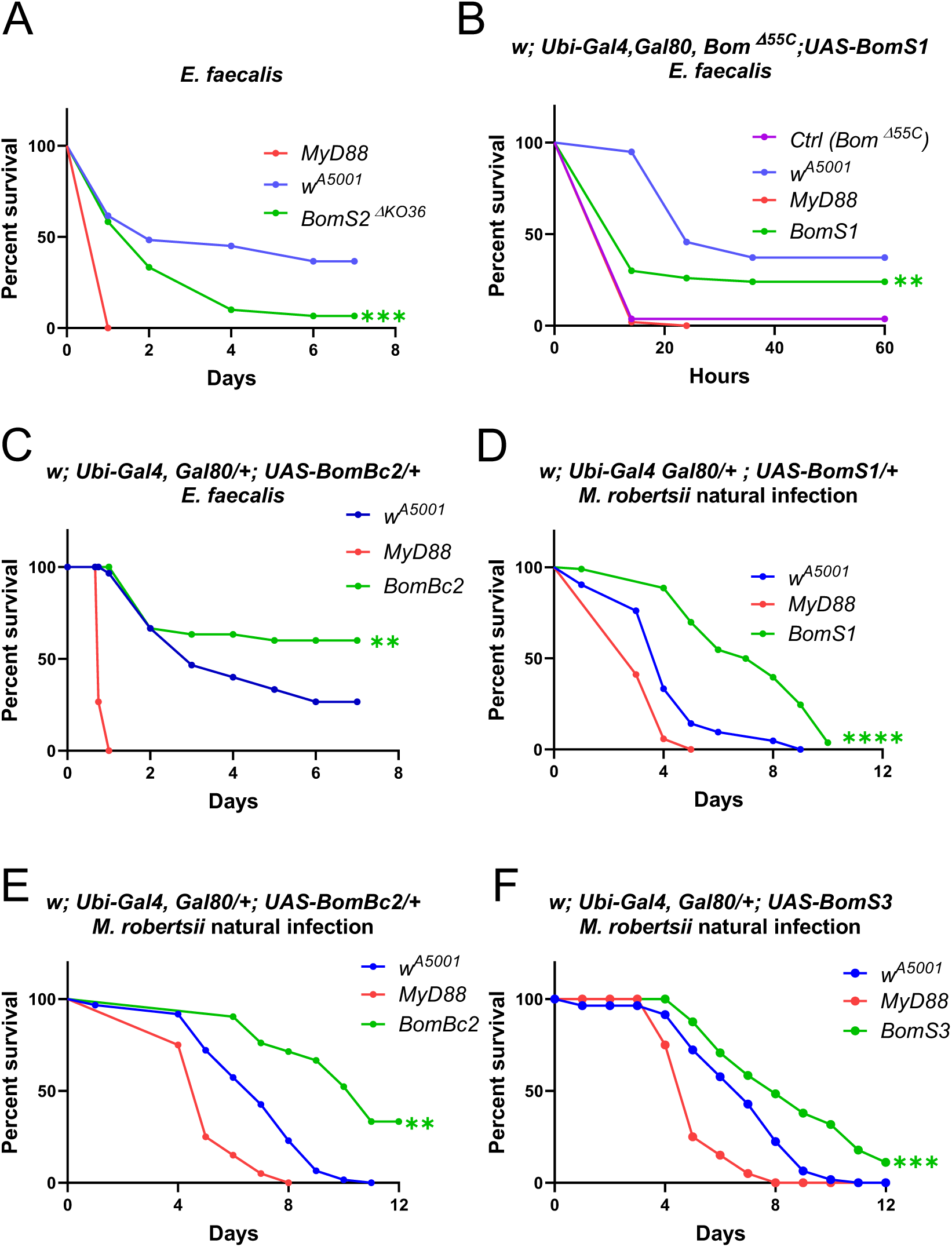
Overexpression of *BomS1, BomBc2, or BomS3* enhances resistance to *E. faecalis* or *M. robertsii*. Complementary to Fig. 2. **(A)** Survival curves of *BomS2^ΔKO36^* null mutant flies after *E. faecalis* infection. **(B)** Survival curves of *BomS1-*overexpressing flies in a *Bom^Δ55C^* background after *E. faecalis* injection. **(C)** Survival experiments of *BomBc2*-overexpression flies in a wild-type background after *E. faecalis* injection. **(D)** Survival curves of *BomS1-*overexpressing flies in a wild-type background against *M. robertsii* natural infection. **(E-F)** Survival curves of *BomBc2* **(E)**, *BomS3* **(F)** overexpressing flies in a wild-type background after *M. robertsii* natural infection. Note: **(A-F)** Each experiment has been performed more than three times. Each experiment used biological triplicates of 20 flies. Pooled data are shown.

**Figure EV4:**
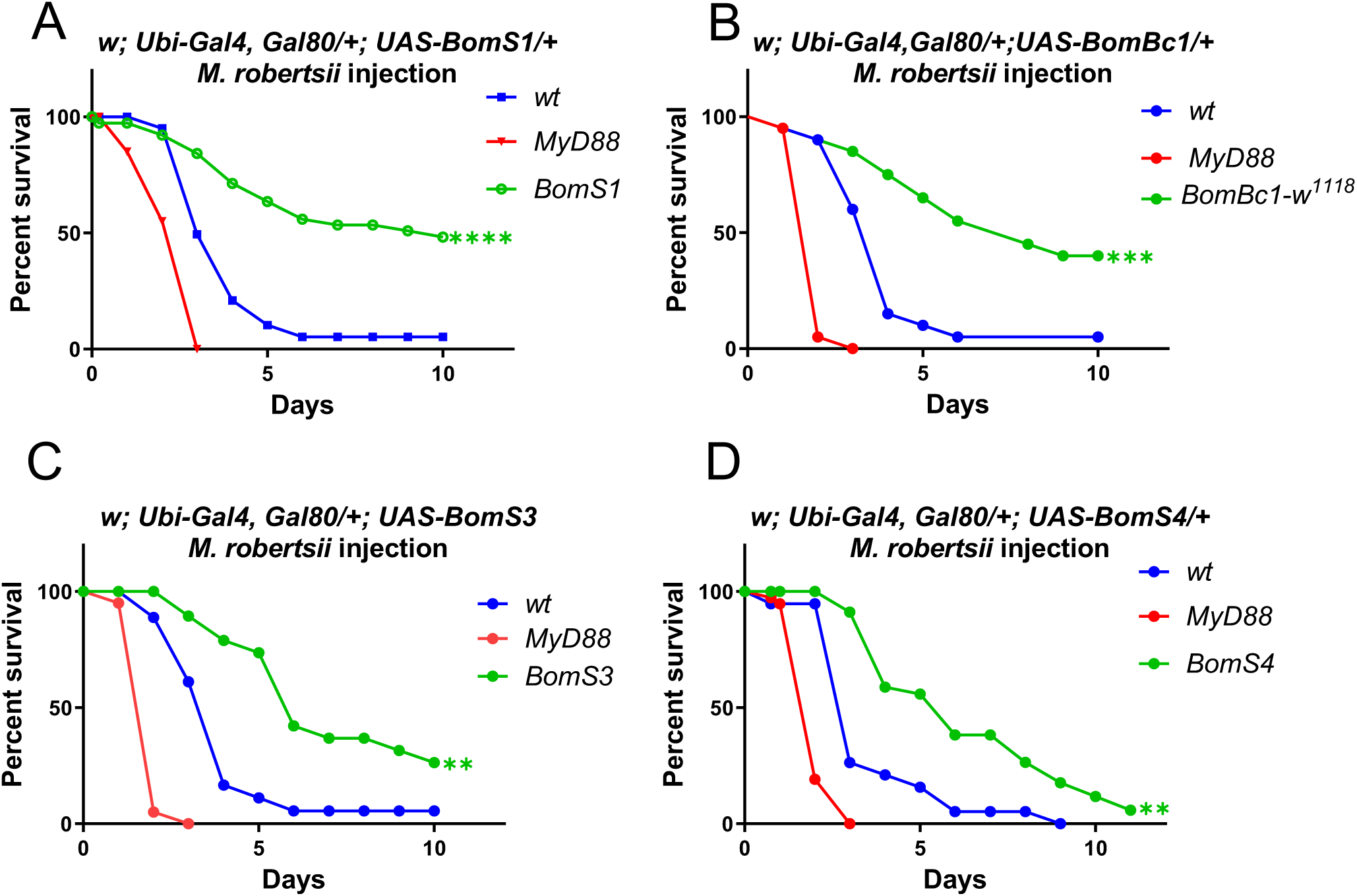
Overexpression of *BomBc1, BomS1, BomS3*, and *BomS4* in a wild type genetic background can protect them from injected *M. robertsii* spores. Complementary to Fig. 3. **(A-D)** Survival curves of *BomBc1* (A), *BomS1* (B), *BomS3* (C), and *BomS4* (D)*-*overexpressing flies in a wild-type background after *M. robertsii* injection. Note: **(A-D)** Each experiment has been performed more than three times. Each experiment used biological triplicates of 20 flies. Pooled data are displayed.

**Figure EV5:**
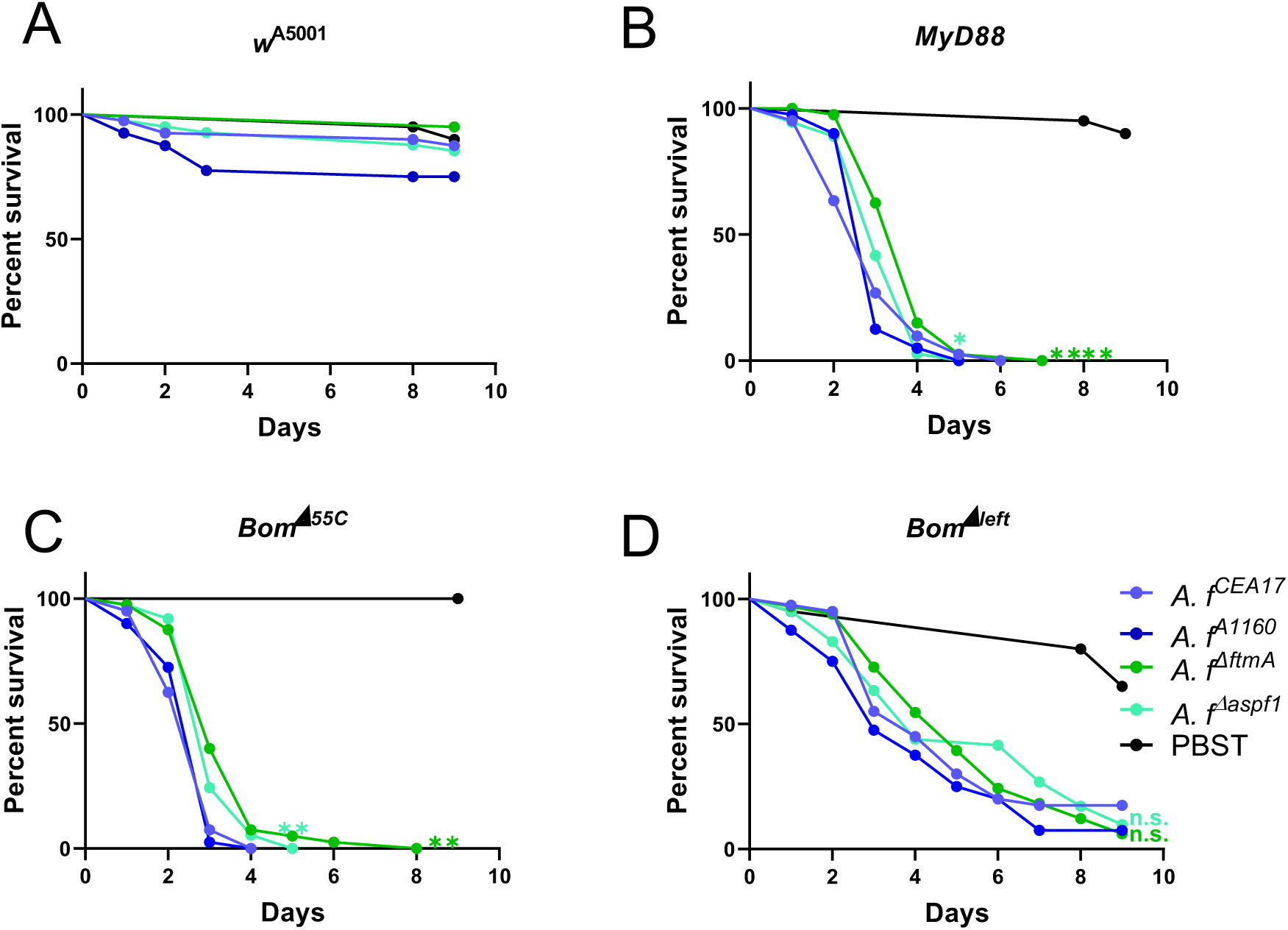
Mycotoxins mutants of *A. fumigatus* are as virulent as wild-type fungi when injected into *Bom^Δleft^* flies. Complementary to Fig. 5. **(A-D)** Survival curves of *w*[A5001] **(A)**, *MyD88* **(B)**, *Bom^Δ55C^* **(C)**, and *Bom^Δleft^* **(D)** flies to *A. fumigatus^ΔftmA^* (fumitremorgin/verruculogen pathway inactivated), *A. fumigatus^Δaspf1^* (restrictocin mutant), and *A. fumigatus* genetic background controls (*A. fumigatus^CEA17^* or *A. fumigatus^A1160^*) injection. Note: **(A-D)** Each experiment has been performed more than three times. Each experiment used biological triplicates of 20 flies. Pooled data are exhibited.

