## Supplementary material and figures for "Distinct Bomanins at the *Drosophila 55C* locus function in resistance and resilience to infections"

**Affiliations:**

[§]Equal contributions

*Corresponding author

### Supplementary material and methods

**Transgenic over-expressed flies in different genetic background:**

**Generation of Barcoded Plasmid Libraries.** To construct barcoded plasmid libraries, we initiated the process by transforming the pGW-HA.attB vector into Escherichia coli DH5α competent cells. Transformed cells were cultured in 500 mL of LB broth and incubated overnight at 37°C with shaking (220 rpm). Plasmid DNA was extracted using a standard alkaline lysis mini-prep protocol. The purified pGW-HA.attB vector was then subjected to double digestion with XhoI and HindIII restriction enzymes (New England Biolabs, Ipswich, MA, USA) according to the manufacturer's instructions.

Barcode oligonucleotides were ligated into the digested vector using T4 DNA ligase (Promega, Madison, WI, USA), following the optimized conditions described in prior work(Bischof, Björklund et al., 2013). Ligation reactions were assessed for efficiency and diversity as previously reported, with the top-performing reactions selected for downstream analysis. On the following day, approximately 600 colonies were screened from ten transformation plates, yielding an estimated library diversity of 60,000 unique barcodes. This diversity metric was calculated by normalizing colony counts to the theoretical transformation efficiency of the *E. coli* strain used.

A barcode diversity exceeding 50-fold the number of intended open reading frame (ORF) clones is recommended to mitigate saturation effects during high-throughput screening. Given our experimental design to over-express 17 target genes, the observed library diversity of 60,000 met this criterion, ensuring adequate representation for our functional genomics studies.

**Generation of Bomanin Transgenic Fly Lines via Gateway Cloning.** 1. Cloning of Bomanin Open Reading Frames (ORFs) into Gateway-Compatible Vectors. The ORFs of Bomanin genes were cloned into the donor vector pDONR™221 (life technologies, no.12536-017) using BP recombination catalyzed by BP Clonase™ II enzyme (Thermo Fisher Scientific) following the manufacturer's protocol. This step generated entry clones for each Bomanin gene. Subsequently, LR recombination, mediated by LR Clonase™ II Plus enzyme (Thermo Fisher Scientific), was employed to transfer the Bomanin ORFs into the pGW-HA.attB-barcoded destination vector, resulting in individual expression vectors for each Bomanin gene. All Gateway reactions were performed according to the manufacturer's optimized conditions, and successful recombination was confirmed by colony PCR using vector-specific primers (**Supplementary Table S1**); 2. Expression Vector Design and Transgenic Fly Generation. Each Bomanin ORF in the destination vector was placed under the control of the UAS-hsp70 promoter to enable tissue-specific expression upon heat shock induction. Transgenic flies were generated by micro-injecting the expression vectors into embryos of the *Drosophila melanogaster* strain BL24749 (Bloomington Drosophila Stock Center), which harbors the attP docking site and ΦC31 integrase on the third chromosome. Injected embryos were reared to adulthood, and F0 males were individually crossed to virgin yw (yellow-white) females. F1 progeny were screened for red-eyed males, indicative of successful transgene integration, using a dissecting microscope (Leica M205 FA, Leica Microsystems, Wetzlar, Germany). Genomic DNA from red-eyed F1 males was subjected to wing-clip PCR with transgene-specific primers (**Supplementary Table S1**) to confirm the presence of the UAS-Bomanin construct; 3. Balancer Crosses and Stock Establishment. Positive F1 males were crossed to virgin females of the third chromosome balancer strain w; TM3/TM6B (Bloomington Drosophila Stock Center). F2 progeny were selected based on red eyes (indicative of the UAS-Bomanin transgene) and the TM6B balancer phenotype(short larvae, Hairy shoulders). Heterozygous F2 males and virgins carrying both the transgene and the balancer were crossed to remove the balancer, generating stable stocks for each Bomanin gene. Transgene integrity was verified by PCR using primers flanking the UAS-Bomanin insertion site , and barcode sequence fidelity was confirmed by Sanger sequencing using vector-specific primers (**Supplementary Table S1**). The original expression vector served as a positive control for all PCR and sequencing reactions.

A detailed list of primers, cloned ORF sequences, and amino acid translations is provided in Supplementary Table S2.

**Overexpression in wild type background.** Following the generation of transgenic UAS fly lines, the initial experimental step involved the collection of virgin females from the ubiquitin promoter-driven Gal4 line (genotype: *w*;*pubi-Gal4*,*ptub-Gal80$^{ts}$*.). These driver virgins were subsequently crossed with males from the transgenic UAS strain at 18°C. Parental flies were transferred to fresh vials every three days to maintain optimal mating conditions and minimize overcrowding effects.

Upon larval hatching, the progeny was shifted to 29°C for a period of 5–7 days to induce Gal80$^{ts}$-mediated derepression of the UAS-driven transgene. Only female offspring were utilized in subsequent infection assays to ensure genetic and physiological uniformity. Overexpression of the target gene (Bomanin) in the wild-type genetic background was confirmed via quantitative reverse transcription PCR (RT-qPCR), with validation data presented in Supplementary Table S5.

**Overexpression in *MyD88* mutant background.** To generate a *MyD88*-deficient genetic background for transgene overexpression, the *w*; *pubi-Gal4*, *ptub-Gal80$^{ts}$* transgenes were recombined onto a second chromosome carrying a MyD88 null allele. Homozygosity of the recombined line was confirmed by two criteria: (1) absence of Drosomycin expression following Enterococcus faecalis infection (as assessed via RT-qPCR or immunoblotting; data not shown) and (2) female sterility. The recombined MyD88-driver line was then crossed to *MyD88/CyO; UAS-Bomanin* flies. Progeny were reared at 18°C and shifted to 29°C for 5–7 days post-hatching to induce Gal80ts-mediated transgene expression. Overexpression of *Bomanin* in the *MyD88* mutant background was validated via RT-qPCR.

**Overexpression in the *Bom$^{\Delta55C}$* deficiency background.** To study Bomanin overexpression in a genomic deficiency background, the *w*;*pubi-Gal4*,*ptub-Gal80$^{ts}$* transgenes were recombined onto a chromosome bearing the *Bom$^{\Delta55C}$* deletion (spanning the Bomanin locus). The recombined line was crossed to *Bom$^{\Delta55C}$/CyO; UAS-Bomanin* flies. Offspring were maintained at 18°C and transferred to 29°C for 5–7 days after hatching to activate transgene expression. Overexpression in the *Bom$^{\Delta55C}$* deletion background was quantified using mass spectrometry (MS) and digital PCR.

**Knock In/out mutant flies:**

**Guide RNA Plasmid Construction.** To achieve high specificity and efficiency in gene targeting, guide RNAs (gRNAs) were designed for each Bomanin-like gene based on the methodology previously reported by Ni and colleagues [178]. Specifically, the protospacer adjacent motif (PAM) sequence (NGG) within the homology arms was removed to minimize off-target effects. The pCFD5 plasmid, a generous gift from the Fillip Port group, served as the backbone for gRNA expression. Following the established protocols from the Fillip Port group, a Gibson assembly cloning kit(Port & Bullock, 2016) was employed to generate plasmids containing two gRNA sequences targeting a single Bomanin gene. Detailed primer sequences used for the construction of gRNA plasmids are provided in the supplementary **Table S3**.

**Donor Plasmid Construction.** The donor plasmid was constructed by first digesting the pBluescript-SK+ plasmid with restriction enzymes PstI and SpeI. The mCherry sequence was obtained from the pUAST-mCherry plasmid, which served as the template for cloning the fluorescent marker. For each gene, the left homology arm (LA) was flanked by a PstI-digested site, while the right homology arm (RA) was flanked by a SpeI-digested site. The donor plasmid was assembled to include the following components: (i) a left arm of approximately 0.8 to 1.0 kb, with the reverse primer positioned immediately upstream of the start codon (ATG); (ii) the mCherry sequence, starting at the ATG and terminating at the stop codon; and (iii) a right arm of approximately 1.0 to 1.5 kb, beginning immediately downstream of the stop codon. Genomic DNA was used as the template for PCR amplification of the homology arms to ensure perfect sequence homology. All assembly reactions were performed using the Gibson assembly cloning kit.

**Microinjection and Generation of Homology Knock-in Flies.** For each gene, the donor plasmid and the corresponding gRNA plasmid were co-injected into lig4 mutant embryos (lig4 line obtained from the Bloomington Drosophila Stock Center, BDSC58492). Following injection, the embryos were allowed to develop into adults, and the resulting F1 progeny were screened for successful integration of the donor plasmid via PCR and sequencing (**Table S4**). To establish stable homology knock-in stocks, the F1 flies were crossed to balancer flies (yw; BcG/CyO). The resulting homozygous or heterozygous knock-in lines were then maintained and characterized for further analysis.

For the knockout strains, *BomS5*$^{\Delta KO}$, *BomS2*$^{\Delta KO6}$, and *BomS2*$^{\Delta KO36}$, we performed isogenization to homogenize their genetic backgrounds, specifically in the wA5001 genetic background.

.

***References***

Bischof J, Björklund M, Furger E, Schertel C, Taipale J, Basler K (2013) A versatile platform for creating a comprehensive UAS-ORFeome library in Drosophila. *Development (Cambridge, England)* 140: 2434-42

Port F, Bullock SL (2016) Augmenting CRISPR applications in Drosophila with tRNA-flanked sgRNAs. *Nature methods* 13: 852-4

***Legends to supplementary figures***

**Figure S1**: **Genetic information of *BomT1*$^{KI}$**

**(A)** Schematic view of the transgenic sequence inserted into the *BomT1* locus to make the *BomT1*$^{KI}$ mutant. The *BomT1* ORF was replaced with the Gal4 sequence, and RFP is used as a fluorescent marker for the presence of the transgene.

**(B)** The transcription level of *Drosomycin* and *Bomanin* genes located at 55C site (except *BomS4*) of *BomT1*$^{KI}$ fly post *M. luteus* injection at 24 hours. This experiment only performed once. The data are presented as means ± SEM.

**FigureS2: Genetic information of *BomBc1*$^{indel}$, and the polymorphism of BomBc1 amino acid sequence in over-expressed transgenic flies**

**(A)** Genomic sequence of the *BomBc1*$^{indel}$ mutant. Notes: The start and stop codons were indicated in red, the same as following in other supplementary figure legends.

**(B)** Amino acid sequence prediction of the *BomBc1*$^{indel}$ mutant. 5 nucleotides(nts) are deleted and give rise to a stop codon, thus only the second Bomanin domain in the C-terminal was removed.

**(C)** The transcription level of *BomBc1*, *Drosomycin* and *BomS1* of *BomBc1*$^{indel}$ post *M. luteus* injection at 24 hours. This experiment only performed once. The data are presented as means ± SEM.

**(D)** Translation sequence of transgenic flies in *BomBc1*$^{w1118}$ and *BomBc1*$^{CS}$ compared with *BomBc1* translation sequence from Flybase (https://flybase.org/). A polymorphism is found in four amino acids which are ‘AQYP’ in Flybase, while ‘TRFS’ in *BomBc1*$^{w1118}$, and ‘TRYP’ in *BomBc1*$^{CS}$.

**Figure S3: Genetic information of the two BomS2 knock out mutants**

*BomS2*$^{\Delta KO6}$, and *BomS2*$^{\Delta KO36}$ were two null mutants. 135nts (counting from ATG) were deleted in *BomS2*$^{\Delta KO6}$, and only 12nts (sequence in blue, GGTGGAAAGTAG) were left in *BomS2*$^{\Delta KO36}$ mutants.

**Figure S4**: **Genetic information of two *BomT2* mutants**

**(A)** *BomT2* knock-in sequence (*BomT2*$^{KI312}$). The sequence underlined and in purple had been replaced by the *mCherry* coding sequence.

**(B)** *BomT2* knock-out sequence (*BomT2*$^{KO259}$). The sequence in green had been deleted. The start and stop codons from the *BomT2* coding sequence (ORF) are indicated in red.

**Figure S5**: **Genetic information of *BomT2*$^{\Delta tail}$**

**(A)** BomT2 sequence in *BomT2*$^{\Delta tail}$ mutant consists on a deletion of 7 nucleotides (TG-AATGA) in the coding sequence (ORF), indicated in green.

**(B)** Amino acid alignment results of *BomT2*$^{\Delta tail}$ flies compared with the *BomT2* wild-type sequence. The mutant predicted sequence formed a stop codon, and the C-terminal tail sequence was deleted. However, the *Bomanin* domain is still present in the mutant line.

**(C)** The expression level of 55C *Bomanin* genes was measured by RT-qPCR in *BomT2*$^{\Delta tail}$ mutants after *M. luteus* injection at 24 hours. This experiment was performed twice. The data are presented as means ± SEM.

**Figure S6**: **Genetic information of *Bom5*$^{\Delta KO}$**

**(A)** The BomS5 mutation *BomS5*$^{\Delta KO}$ consists in a 153-nucleotides deletion inside the coding sequence (ORF), indicated in green.

**(B)** Translation sequence alignment of *BomS5*$^{\Delta KO}$. Peptide sequence alignment between *BomS5*$^{\Delta KO}$ and the wild-type BomS5.

**(C)** Expression level of other 55C locus *Bom* genes measured by RT-qPCR in *BomS5*$^{\Delta KO}$ mutants after *M. luteus* injection at 24 hours. Each experiment has been performed more than three times. The data are presented as means ± SEM.

**Figure S7: The efficiency of knock down flies of *BomT1, BomBc1,* and *BomBc2.***

**(A-D)** Knock-down efficiency of *BomT1* **(A)**, two *BomBc1* **(B)**, and two *BomBc2* **(C, D)** knock-down flies measured by RT-qPCR. Those experiment only performed once. The data are presented as means ± SEM and analyzed using Kruskal-Wallis test; *** $p<0.001$.

**Figure S8**: **The efficiency of overexpression flies of single *Bomanin* genes in wild-type background.**

Expression level of single *Bomanin* gene overexpressing flies in a wild-type background by RT-qPCR. The genotype of transgenic flies: *w; Ubi-Gal4, Gal80ts/+; UAS-Bom/+*. Each experiment has been performed at least three times. The data are presented as means ± SEM and analyzed using ANOVA (one-way) with Dunnett's multiple comparisons test. * p<0.05; ** p<0.01; *** p<0.001; **** p<0.0001; ns, no significant difference.

**Figure S9**: **The efficiency of overexpression flies of single *Bomanin* genes in *Bom*$^{\Delta 55C}$ background.**

**(A)** The translation level of single *BomS*-overexpressing flies in the *Bom*$^{\Delta 55C}$ background measured by MALDI-TOF mass spectrometry. More than 5 flies were tested for each *BomS*.

(B) Expression level of *BomTs* and *BomBcs* overexpressing flies in the *Bom*$^{\Delta 55C}$ background by RT-qPCR. This experiment only performed once. The data are presented as means ± SEM.

**Figure S10: Differential protection and genetic background dependence of the *Bomanin* genes in *Drosophila* against *E.faecalis* and *C. albicans* infections**

**(A)** Survival curves of the *BomT1*-overexpressing flies in a wild-type background after *E. faecalis* infection.

**(B-C)** Survival curves of *BomBc1*$^{V15384}$ **(B)** and *BomBc1*$^{BL65901}$ **(C)** line after *C. albicans* septic injury.

**(D-E)** Survival curves of *BomT2*$^{\Delta tail}$ **(D)** and two *BomT2* null mutants **(E)** after *C. albicans* infection.

**(F)** Survival curves of *BomBc1*$^{w1118}$ isoform overexpressing flies in a *Bom*$^{\Delta 55C}$ background after *C. albicans* infection.

Note: **(A-F)** Each experiment has been performed at least three times. Each experiment used biological triplicates of 20 flies. The data were analyzed using Log-Rank test. ** p<0.01; **** p<0.0001; ns, no significant difference.

**Figure S11**: **The phenotype of *BomS1*-overexpressing flies in a *MyD88* mutant background**

**(A)** Survival curves of *BomS1*-overexpressing flies in a *MyD88* mutant background after *C. glabrata* infection. The experiment has been performed more than three times. Each experiment used biological triplicates of 20 flies. The data of were analyzed by using Log-Rank test; ** p<0.01.

**(B-D)** The bristles of *BomS1*-overexpressing flies in a wild-type background. Bristle of a wild-type fly **(B)**. Bristle from the TM3Sb balancer phenotype **(C)**. Bristle of a *BomS1*-overexpressing fly in a wild-type background **(D)**.

**Figure S12**: **Biosynthesis of Fumitremorgins/verruculogen in *A. fumigatus*. https://enzyme-database.org/reaction/alkaloid/fumitrem2.html**

**Figure S13**: **Susceptibility of *Bombardier* (*Bbd*) mutants to *A. fumigatus* conidia or mycotoxin injections.**

**(A-D)** Survival curves of the *Bombardier* mutants after *A. fumigatus* **(A)**, verruculogen **(B),** Restrictocin **(C)**, and gliotoxin **(D)** injection. Each experiment has been performed at least three times. Each experiment used a biological repetition of 20 flies.

### Figure S1

A

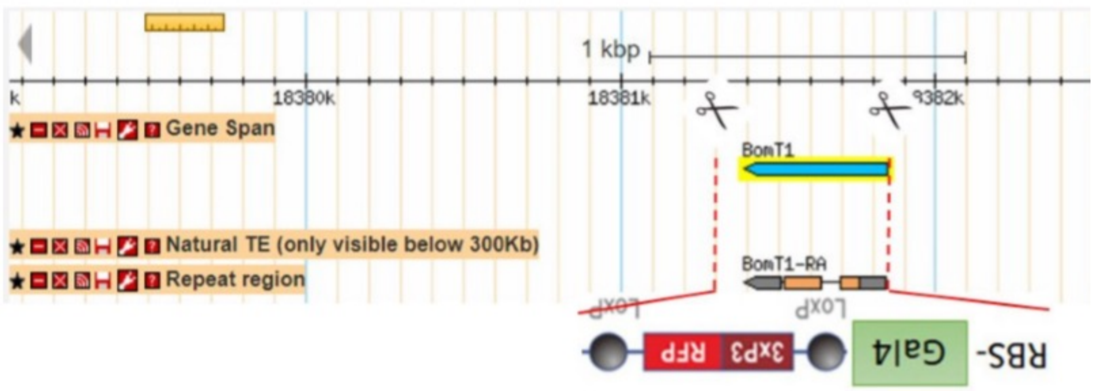
Genome Editing:
1 kbp
18380k
18381k
9382k
Gene Span
Natural TE (only visible below 300Kb)
Repeat region
BomT1
BomT1-RA
LoxP
LoxP
RFP
3xP3
Gal4
RBS-

B

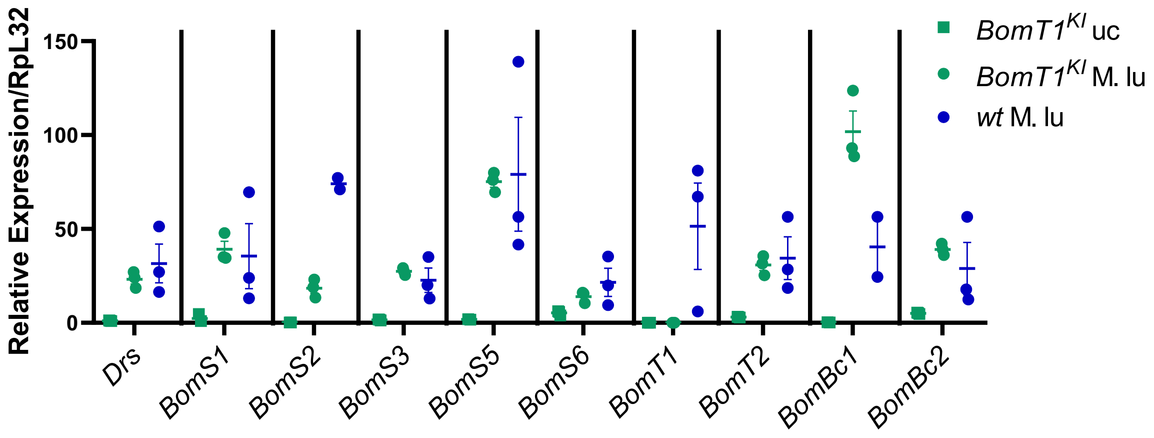
Relative Expression/RpL32
150
100
50
0
Drs
BomS1
BomS2
BomS3
BomS5
BomS6
BomT1
BomT2
BomBc1
BomBc2
BomT1KI uc
BomT1KI M. lu
wt M. lu

### Figure S2

**A**

*BomBc1* *Indel*

BomBc1>2R:18382704..18383235 (reverse complemented)

```
TCGAATAGATTCGTCTTGCACGCAGATTGAGAATGAAGTGCCTGATTCTGTCCTTTGCAATTTTCG
TTGTCCTGGCTTCCCAGGCTACGGgtaagaatcagatgggcgcgcctttcaatgtcacaatttcgc
ttattattttattattccgcagCCGGAAATGTGATTATCGGCGGAGTATGCCAGGATTGCAGTCCG
CCGGTGGCGGAAAACGTCGTAGTCGGTGGCCAATCCTACAGGACGGGTAGGCCGGGCCAGGGAACG
GTGTATATCAATTCCCCTGGCGCATATCTAGGAGCTCTCGATGGTCCCATTCGGCGAACTGGCGCT
GGCGGCGGAGGAGGCGGTGGCGCCCAGTATCCGGATGGTTACAGTGGTCGTCTGCCAGGTGGCACT
TACCTTCACAATAAGGATTGCGTGGGCTGCAGCATCAGCGGGGGCGGGGATTAACGAAATTATATT
GTGATTTGTACATAAAATACCTCGCATTTTTTAATTTTCGGGCATAATTTATAATTCTTTAAAAAA
TCTG
```

Deleted seq: **XXXXX**
Formed stop codon: **TAG**
*BomBc1* ORF: **ATG...TAA**

**B**

```
                                                         N-Bomanin domain                  Stop codon
BomBc1_indel_mutant_aa.seq MKCLILSFAIFVVLASQATAGNVIIGGVCQDCSPPVAENVVVGGQSYRTGRPGQGTVY--
BomBc1_aa.seq              MKCLILSFAIFVVLASQATAGNVIIGGVCQDCSPPVAENVVVGGQSYGAG-PGNG-VYQF
                           *********************************************** .* **.* **

                                                                               C-Bomanin domain
BomBc1_indel_mutant_aa.seq ---INSPGAYLGALDGPIRRTGAGGGGGGGAQYPDGYSGRLPGGTYLHNKDCVGCSISGGGD
BomBc1_aa.seq              PWRISRSSRWSHSANWRWRRRRRWRPVSGWLQWSSAR-WHLPSQGLRGLQHQRGRGL---
                           *         .      **         *   *       .**       .    *   .
```

**C**

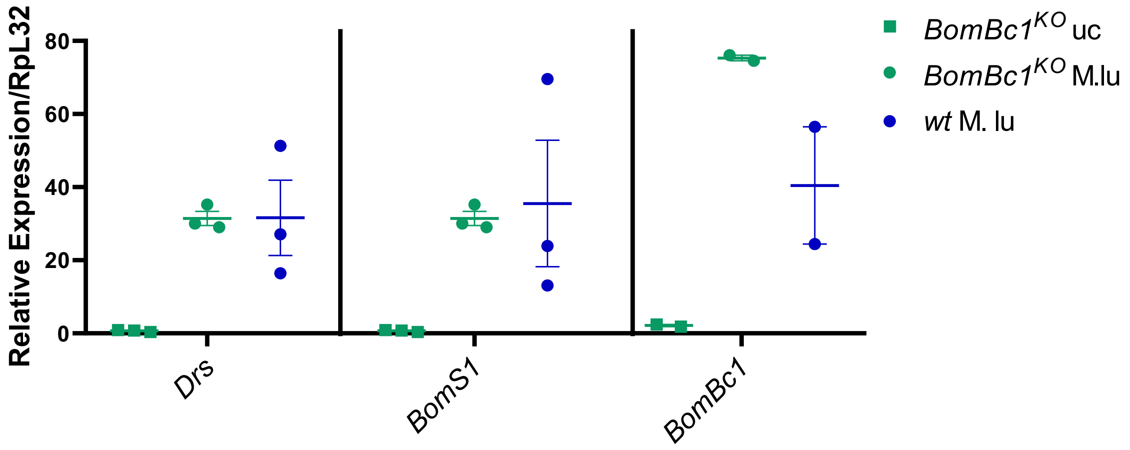

**D**

```
                                           Bomanin domain
BomBc1_aa.seq      MKCLILSFAIFVVLASQATAGNVIIGGVCQDCSPPVAENVVVGGQSYRTGRPGQGTVYIN
BomBc1-CS_aa.seq   MKRLILSFAIFVVLASQATAGNVIIGGVCQDCSPPVAENVVVGGQSYRTGRPGQGTVYIN
BomBc1-W_aa.seq    MKCLILSFAIFVVLASQATAGNVIIGGVCQDCSPPVAENVVVGGQSYRTGRPGQGTVYIN
                   ** *********************************************************

                                           Polymorphisms        Bomanin domain
BomBc1_aa.seq      SPGAYLGALDGPIRRTGAGGGGGGAQYPDGYSGRLPGGTYLHNKDCVGCSISGGGD
BomBc1-CS_aa.seq   SPGAYPGALDGPIRRTGAGGGGGGTRYPDGYSGRLPGGTYLHNKDCVGCSISGGGD
BomBc1-W_aa.seq    SPGAYPGALDGPIRRTGAGGGGGGTRFSDGYSGRLPGGTYLYNKDCVGCSISGGGD
                   ***** *******************...*************** ****************
```

### Figure S3

BomS2(IM2, CG18106) >2R:18386602..18387030
CCAAGAGCATCAGTTGAATTCAATCGATTGCTTGTGCATTTAGCAAATCAAAGCCACAACAACC
AAACCAGAATCAATATGAAGTTCTTCTCAGTCGTCACCGTCTTTGTGTTCGGTCTGCTGGCTCT
GGCCAACGgttagtaatactatattttatagctatttgattactttataaattatttatttctt
ccgcagCTGTTCCCCTGTCGCCCGATCCAGGAAATGTGGTAATCAACGGGGACTGCAAATACTG
CAATGTGCACGGTGGAAAGTAGGAAAGTAGGAAAGTAGGAAAGTACTCGCCTTAATTTCGAAGA
TGGGCCAAAACTTACCTCAAATCCAAAGCACCATATTTATACTCTCACTCTTGTACTAAAATGA
AAACTAGTAGTAAAAAATACATCGCCAATTACAAATAAAATGGTGAAAAAAC

*BomS2* $^{\Delta KO6}$ knock out sequence

*BomS2* $^{\Delta KO36}$ knock out sequence

*BomS2* ORF ATG…GGTGGAAAGTAG

Intron sequence: gttagtaatactatattttatagctatttgattactttataaattatttatttcttccgcag

### Figure S4

**A**

*BomT2*$^{KI312}$

>2R:18388995..18389443

CAATTATATATATAATATAGCAAATACTATATGTACTATGGCATATAAAAAGCAGAGCACTCGCACTTT
GGCAGTCAGATTTATTCGAAATCCCCAAGGAAAAAC**ATG**AAAGCTCTTCAAGTCGCCGGAACTTTGATG
CTGCTTTTCTGCCTGCTGGCAGCTGTTAATGgtaattatttaatgaaataataaaatatatatatgata
tttttagctaatcatttccttattgcagCTACGCCGGGACAAGTGTATATCAATGGGAAATGCATTGAC
TGCAATAAGCCTGATAATGATCCGGGAATTATAATTCCTCCAGACCATAAATCAGCTGGATCCATGTCT
TACACACTCACATCTGGAGCCATCTTCTTTGGAATTATATATCATATATTCAGT**TAA**ATTACTATGTAA
TCAATAAGTTAATAAAATAATATCTTTTATTTCTCaaaaaaaaaccttaagcttgtgacaaattgcgaa
attgcgaa

**Replaced sequence: XXXXXX**
***BomT2* ORF: ATG…TAA**
**Intron sequence:**
gtaattatttaatgaaataataaaatatatatatgatatttttagctaatcatttccttattgcag

**B**

*BomT2*$^{KO259}$

>2R:18388995..18389443

CAATTATATATATAATATAGCAAATACTATATGTACTATGGCATATAAAAAGCAGAGCACTCGCACTTT
GGCAGTCAGATTTATTCGAAATCCCCAAGGAAAAAC**ATG**AAAGCTCTTCAAGTCGCCGGAACTTTGATG
CTGCTTTTCTGCCTGCTGGCAGCTGTTAATGgtaattatttaatgaaataataaaatatatatatgata
tttttagctaatcatttccttattgcagCTACGCCGGGACAAGTGTATATCAATGGGAAATGCATTGAC
TGCAATAAGCCTGATAATGATCCGGGAATTATAATTCCTCCAGACCATAAATCAGCTGGATCCATGTCT
TACACACTCACATCTGGAGCCATCTTCTTTGGAATTATATATCATATATTCAGT**TAA**ATTACTATGTAA
TCAATAAGTTAATAAAATAATATCTTTTATTTCTCaaaaaaaaaccttaagcttgtgacaaattgcgaa
attgcgaa

**Knock out sequence: XXXXXX**
***BomT2* ORF: ATG…TAA**

### Figure S5

## A

**BomT2 (IM28, CG16836)** >2R:18388995..18389443

*BomT2$^{\Delta tail}$*

```
TAATTATATATATATAATATAGCAAATACTATATGTACTATGGCATATAAAAAGCAGAGCACTCGC
ACTTTGGCAGTCAGATTTATTCGAAATCCCCAAGGAAAAACATGAAAGCTCTTCAAGTCGCCGG
AACTTTGATGCTGCTTTTCTGCCTGCTGGCAGCTGTTAATGgtaattatttaatgaaataataa
aatatatatatgatatttttagctaatcatttccttattgcagCTACGCCGGGACAAGTGTATA
TCAATGGGAAATGCATTGACTGCAATAAGCCTGATAATGATCCGGGAATTATAATTCCTCCAGA
CCATAAATCAGCTGGATCCATGTCTTACACACTCACATCTGGAGCCATCTTCTTTGGAATTATA
TATCATATATTCAGTTAAATTACTATGTAATCAATAAGTTAATAAAATAATATCTTTTATTTCT
C
```

**knock out nucleotides : TGATAATGA**

***BomT2* ORF: ATG…TAA**

## B

```
                                                   Bomanin domain         Stop codon
BomT2.seq                 MKALQVAGTLMLLFCLLAAVNATPGQVYINGKCIDCNKPDNDPGIIIPPDHKSAGSMSYT
BomT2_Tail_mutant.seq     MKALQVAGTLMLLFCLLAAVNATPGQVYINGKCIDCNKPFRE--LFLQTINQLDPCLTHS
                          ****************************************   .  . .    ..    ... .

BomT2.seq                 -LTSGAIFFGIIYHIFS
BomT2_Tail_mutant.seq     HLEPSSLELYIIYSV--
                           *   ..   *** .
```

## C

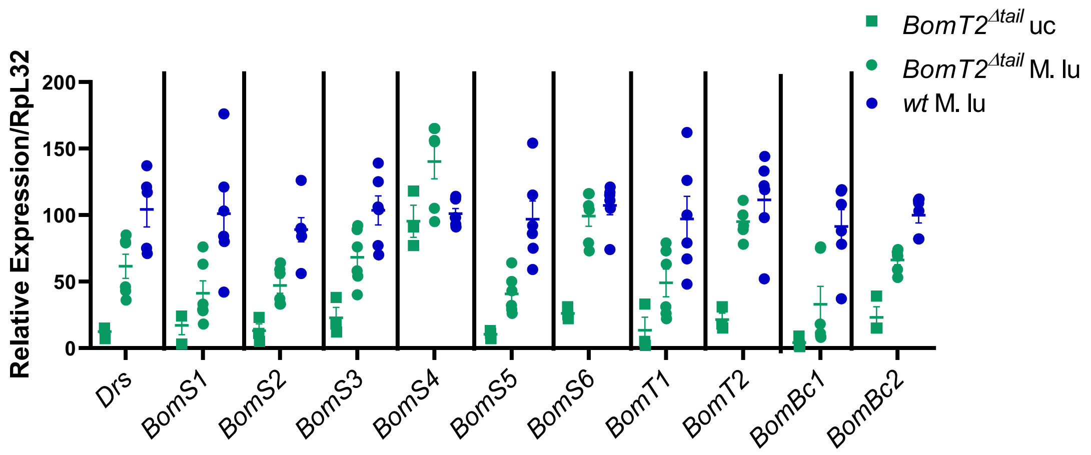

### Figure S6

**A**

***BomS5$^{\Delta KO}$***

>2R:18389959..18390242

ATCAGTTTCATTTCAACCGTTGCCAACAAT**ATG**AAGTGGATGTCCTTGGTCTTTCTATGCGGTCTGC
TCGCCATGGCAGTGGgtgagtatctataagattcatacatccttaaccagtatcctaattatttttc
tactattactatagCTTCTCCGTTAAATCCGGGTAATGTCATTATCAATGGAGATTGCCGTCATTGT
AATGTTCGCGGAGGC**TAA**ATTGGAGTTATAAAACTTTGATGTTTAGTAACTACAATAAAACTTATGA
AAAATACAGATTGTGC

**Knock out sequence: XXXXXX**
***BomS5* ORF: ATG…TAA**
**Intron sequence:**
gtgagtatctataagattcatacatccttaaccagtatcctaattatttttctactattactatag

**B**

```
                                              Bomanin domain
BomS5_KO_mutant.seq MKWMSLVFLCG-----------------------------
BomS5.seq           MKWMSLVFLCGLLAMAVASPLNPGNVIINGDCRHCNVRGG
                    ***********
```

**C**

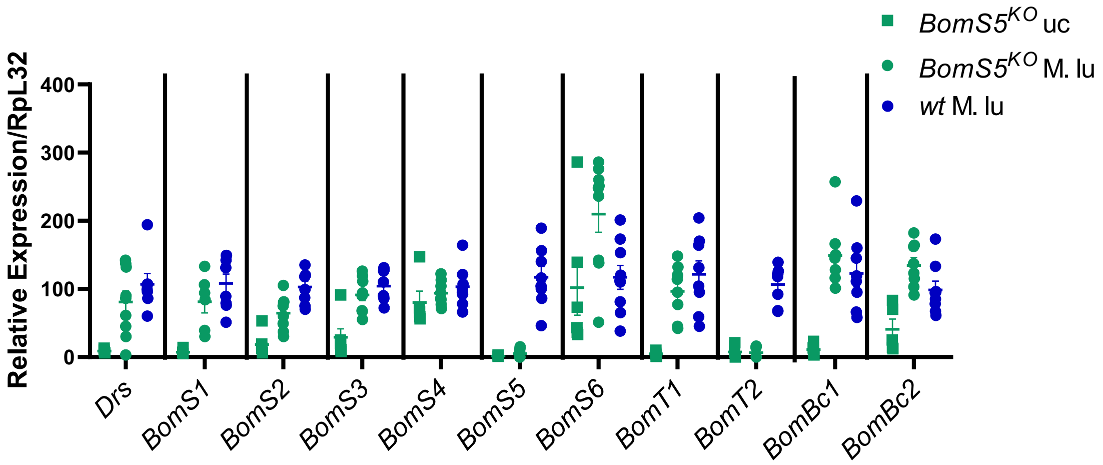

### Figure S7

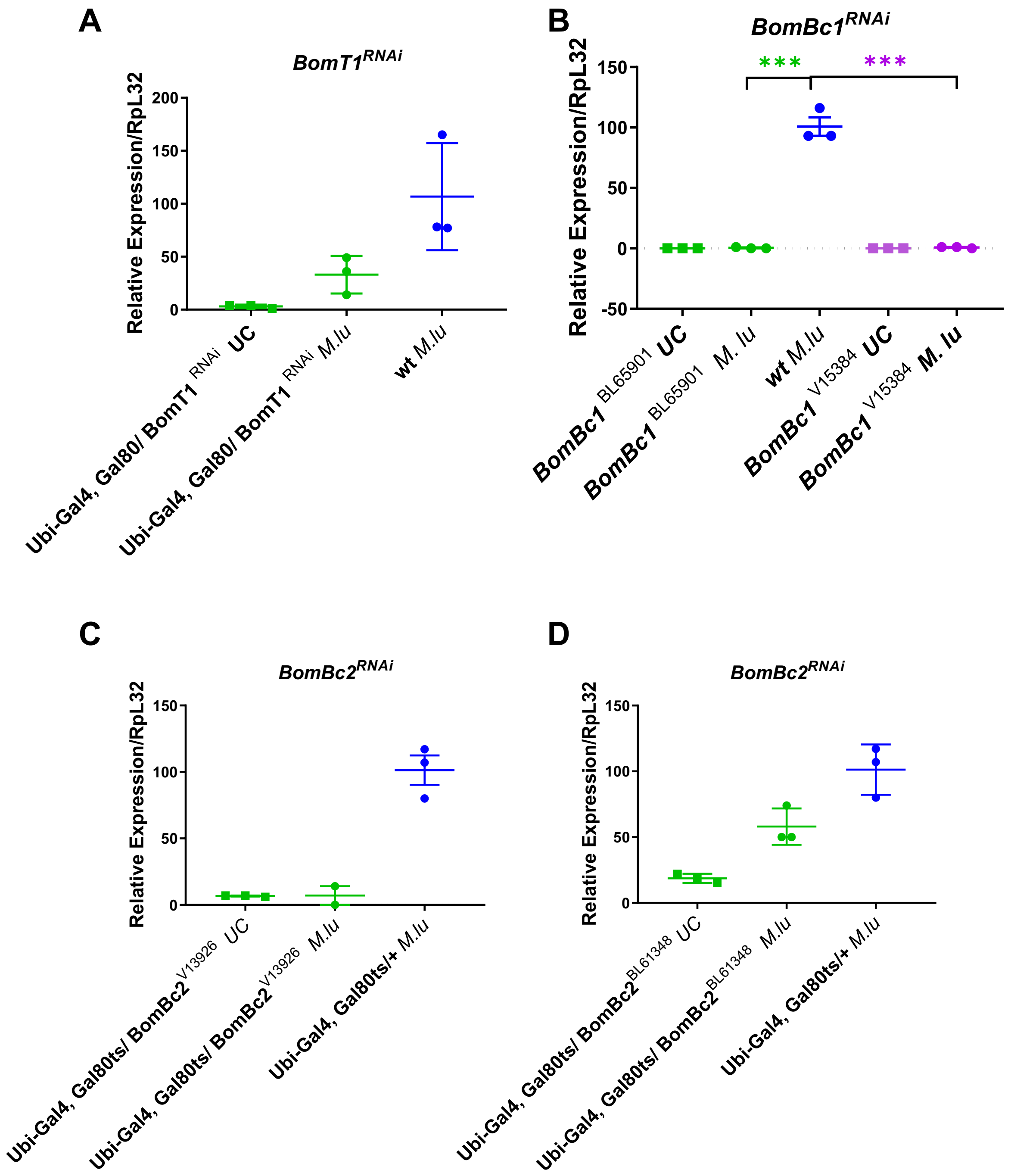

A
BomT1RNAi
Relative Expression/RpL32
200
150
100
50
0
Ubi-Gal4, Gal80/ BomT1 RNAi UC
Ubi-Gal4, Gal80/ BomT1 RNAi M.lu
wt M.lu
B
BomBc1RNAi
***
***
Relative Expression/RpL32
150
100
50
0
-50
BomBc1 BL65901 UC
BomBc1 BL65901 M. lu
wt M.lu
BomBc1 V15384 UC
BomBc1 V15384 M. lu
C
BomBc2RNAi
Relative Expression/RpL32
150
100
50
0
Ubi-Gal4, Gal80ts/ BomBc2V13926 UC
Ubi-Gal4, Gal80ts/ BomBc2V13926 M.lu
Ubi-Gal4, Gal80ts/+ M.lu
D
BomBc2RNAi
Relative Expression/RpL32
150
100
50
0
Ubi-Gal4, Gal80ts/ BomBc2BL61348 UC
Ubi-Gal4, Gal80ts/ BomBc2BL61348 M.lu
Ubi-Gal4, Gal80ts/+ M.lu

### Figure S8

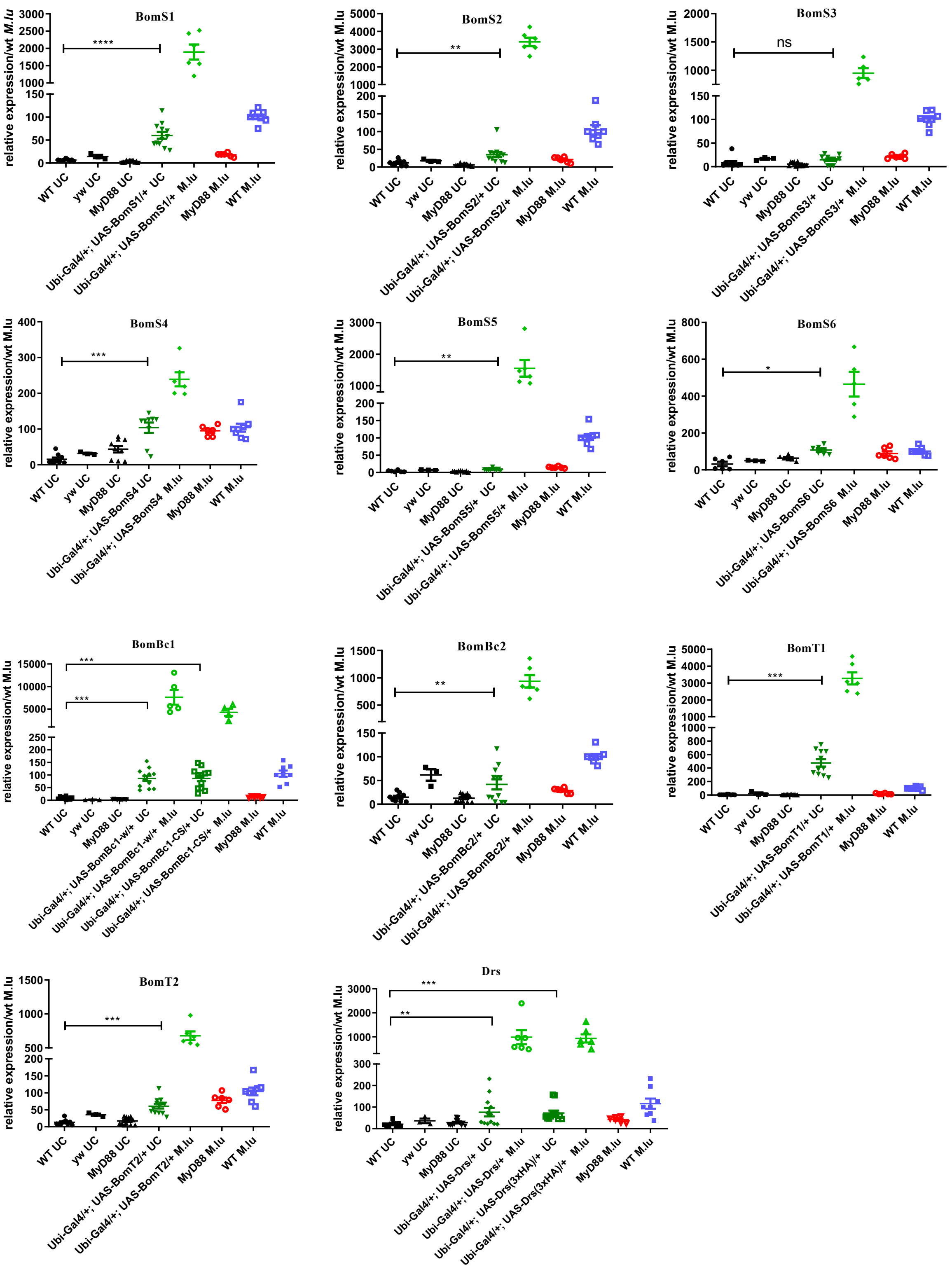

BomS1
BomS2
BomS3
BomS4
BomS5
BomS6
BomBc1
BomBc2
BomT1
BomT2
Drs
relative expression/wt M.lu
****
**
ns
***
*
WT UC
yw UC
MyD88 UC
MyD88 M.lu
WT M.lu
Ubi-Gal4/+; UAS-BomS1/+ UC
Ubi-Gal4/+; UAS-BomS1/+ M.lu
Ubi-Gal4/+; UAS-BomS2/+ UC
Ubi-Gal4/+; UAS-BomS2/+ M.lu
Ubi-Gal4/+; UAS-BomS3/+ UC
Ubi-Gal4/+; UAS-BomS3/+ M.lu
Ubi-Gal4/+; UAS-BomS4 UC
Ubi-Gal4/+; UAS-BomS4 M.lu
Ubi-Gal4/+; UAS-BomS5/+ UC
Ubi-Gal4/+; UAS-BomS5/+ M.lu
Ubi-Gal4/+; UAS-BomS6 UC
Ubi-Gal4/+; UAS-BomS6 M.lu
Ubi-Gal4/+; UAS-BomBc1-w/+ UC
Ubi-Gal4/+; UAS-BomBc1-w/+ M.lu
Ubi-Gal4/+; UAS-BomBc1-CS/+ UC
Ubi-Gal4/+; UAS-BomBc1-CS/+ M.lu
Ubi-Gal4/+; UAS-BomBc2/+ UC
Ubi-Gal4/+; UAS-BomBc2/+ M.lu
Ubi-Gal4/+; UAS-BomT1/+ UC
Ubi-Gal4/+; UAS-BomT1/+ M.lu
Ubi-Gal4/+; UAS-BomT2/+ UC
Ubi-Gal4/+; UAS-BomT2/+ M.lu
Ubi-Gal4/+; UAS-Drs/+ UC
Ubi-Gal4/+; UAS-Drs/+ M.lu
Ubi-Gal4/+; UAS-Drs(3xHA)/+ UC
Ubi-Gal4/+; UAS-Drs(3xHA)/+ M.lu

### Figure S9

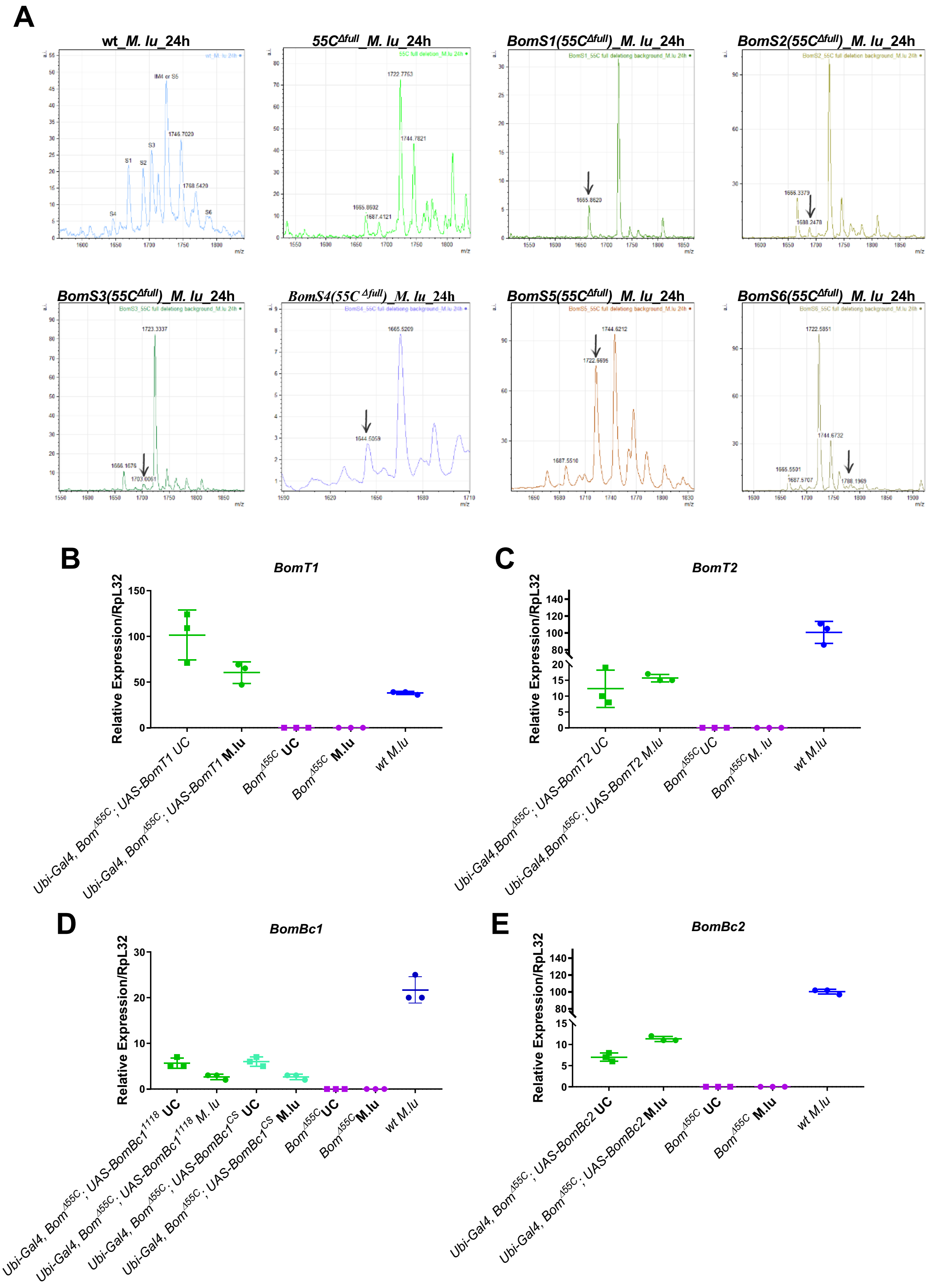

A
wt_M. lu_24h
55CΔfull_M. lu_24h
BomS1(55CΔfull)_M. lu_24h
BomS2(55CΔfull)_M. lu_24h
BomS3(55CΔfull)_M. lu_24h
BomS4(55CΔfull)_M. lu_24h
BomS5(55CΔfull)_M. lu_24h
BomS6(55CΔfull)_M. lu_24h
B
BomT1
C
BomT2
D
BomBc1
E
BomBc2
Relative Expression/RpL32
wt M.lu

### Figure S10

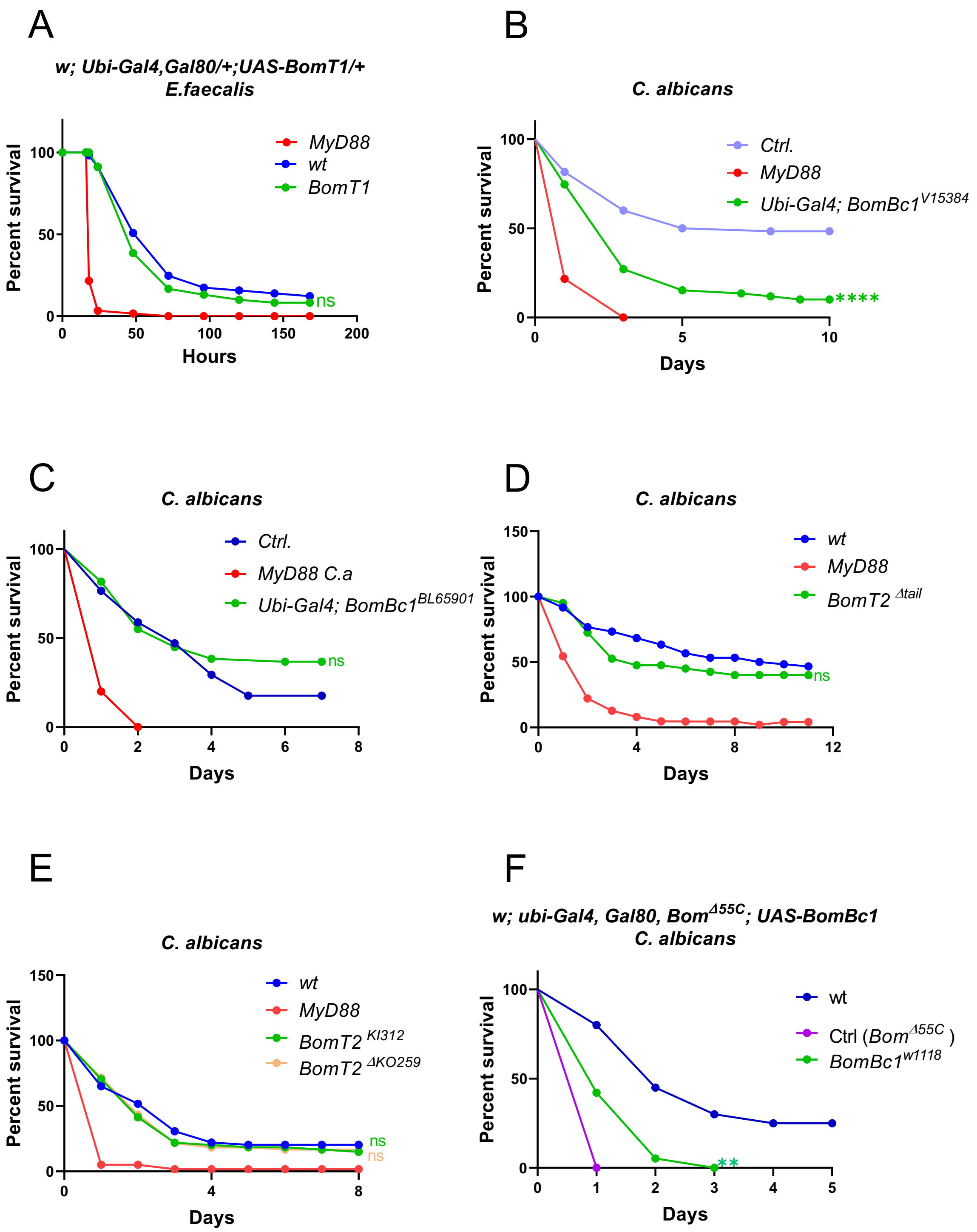

A
w; Ubi-Gal4,Gal80/+;UAS-BomT1/+
E.faecalis
Percent survival
Hours
MyD88
wt
BomT1
ns
B
C. albicans
Percent survival
Days
Ctrl.
MyD88
Ubi-Gal4; BomBc1V15384
****
C
C. albicans
Percent survival
Days
Ctrl.
MyD88 C.a
Ubi-Gal4; BomBc1BL65901
ns
D
C. albicans
Percent survival
Days
wt
MyD88
BomT2 Δtail
ns
E
C. albicans
Percent survival
Days
wt
MyD88
BomT2 KI312
BomT2 ΔKO259
ns
ns
F
w; ubi-Gal4, Gal80, BomΔ55C; UAS-BomBc1
C. albicans
Percent survival
Days
wt
Ctrl (BomΔ55C )
BomBc1w1118
**

**Figure S11**

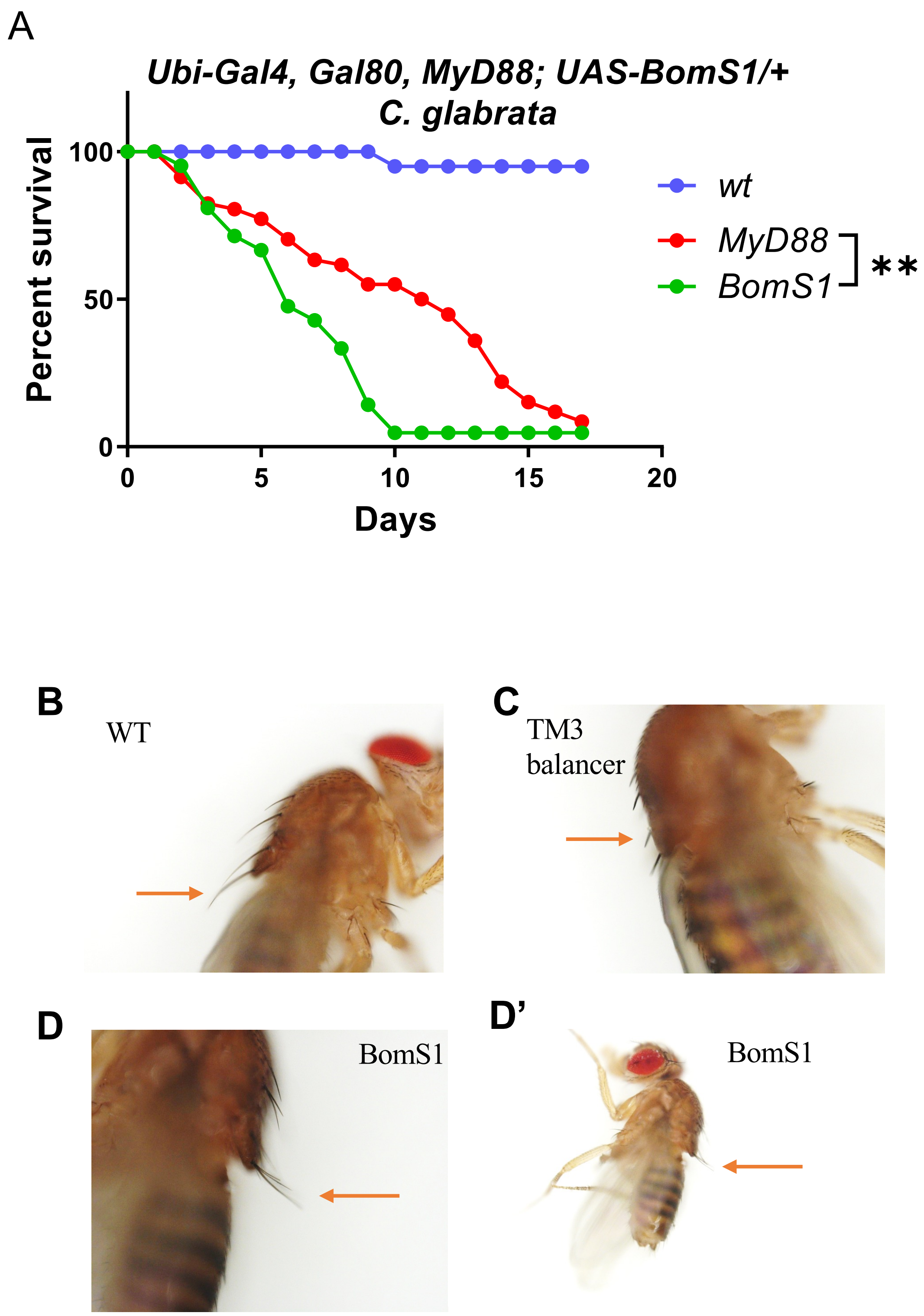
A
Ubi-Gal4, Gal80, MyD88; UAS-BomS1/+
C. glabrata
Percent survival
100
50
0
0
5
10
15
20
Days
wt
MyD88
BomS1
**
B
WT
C
TM3
balancer
D
BomS1
D'
BomS1

### Figure S12

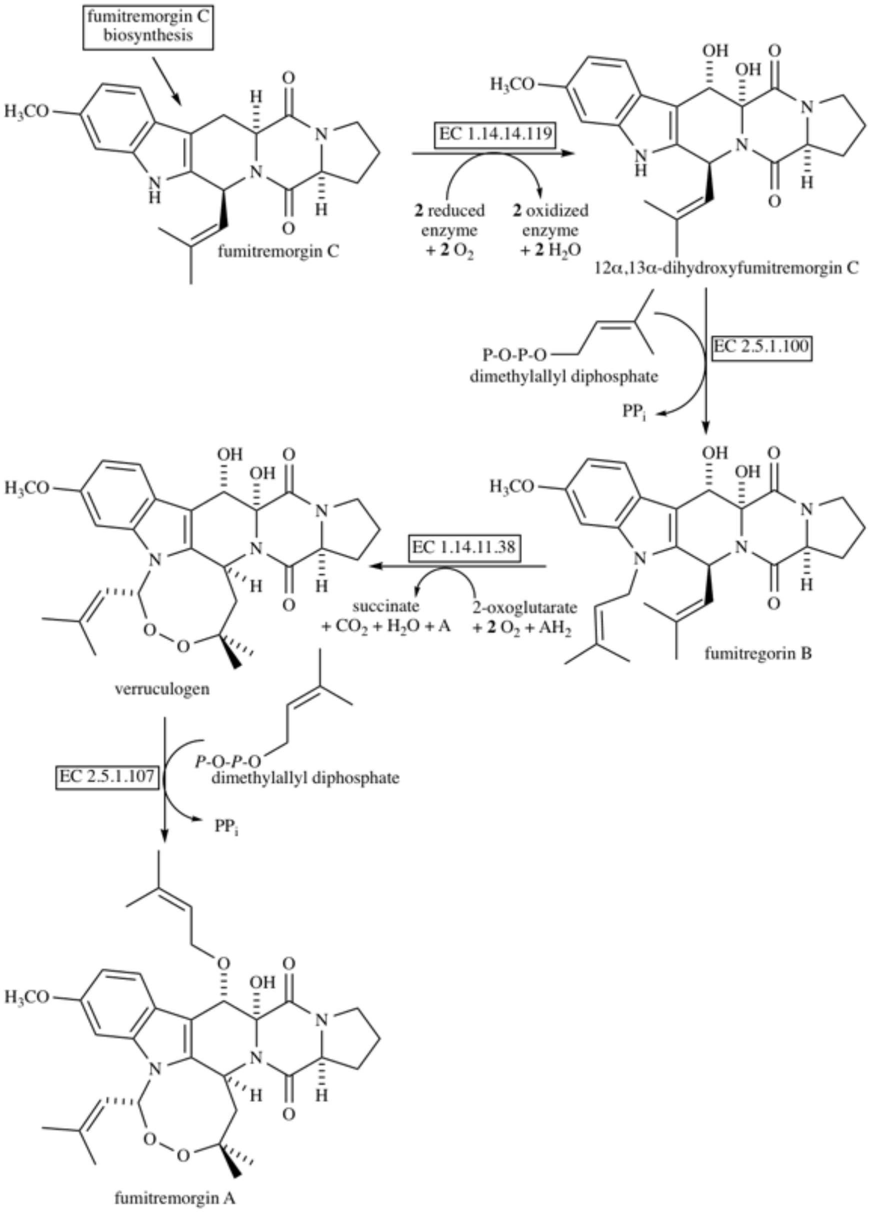
fumitremorgin C biosynthesis
H3CO
fumitremorgin C
EC 1.14.14.119
2 reduced enzyme + 2 O2
2 oxidized enzyme + 2 H2O
12α,13α-dihydroxyfumitremorgin C
EC 2.5.1.100
P-O-P-O
dimethylallyl diphosphate
PPi
fumitregorin B
EC 1.14.11.38
succinate + CO2 + H2O + A
2-oxoglutarate + 2 O2 + AH2
verruculogen
EC 2.5.1.107
P-O-P-O
dimethylallyl diphosphate
PPi
fumitremorgin A

### Figure S13

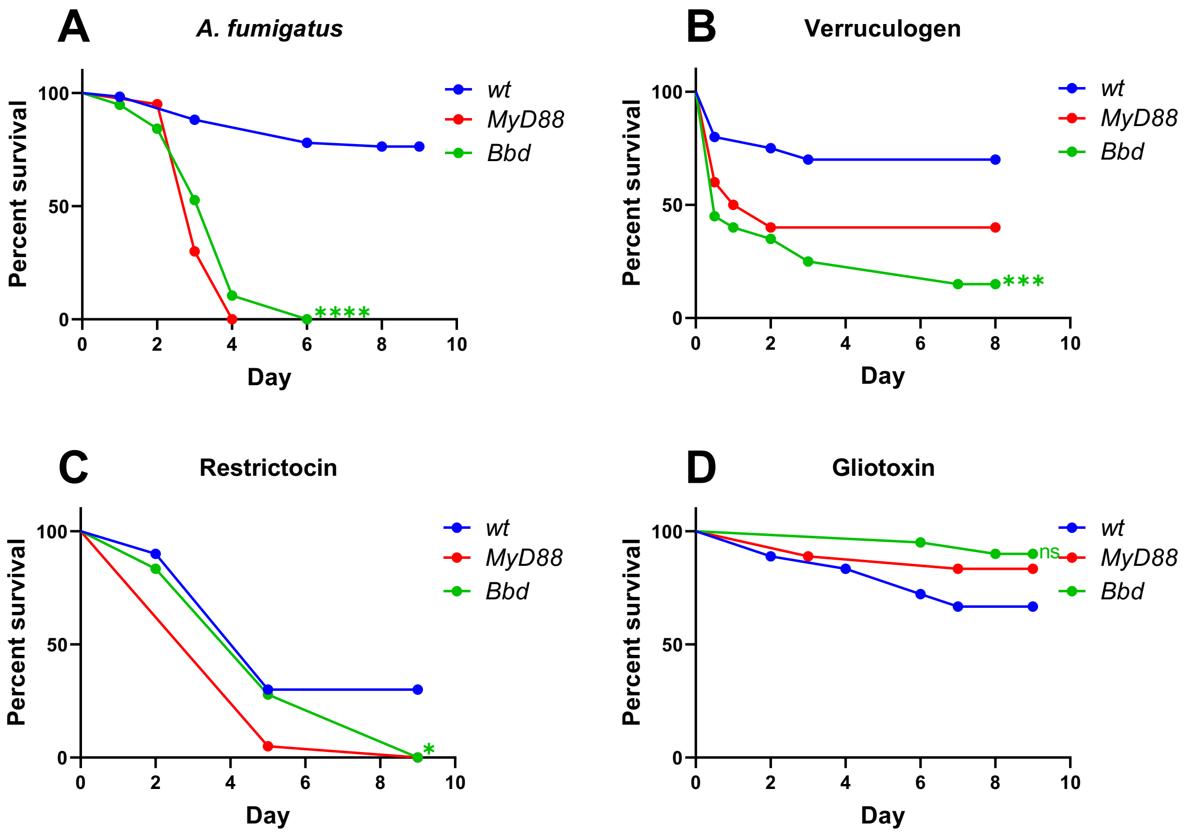

A
A. fumigatus
Percent survival
100
50
0
Day
0
2
4
6
8
10
wt
MyD88
Bbd
****
B
Verruculogen
***
C
Restrictocin
*
D
Gliotoxin
ns

**Table S1: synthesized primer sequence for expression vector.**

| Expression vector clone primer<br>primer sequence | sequence name | Bomanin |
|---|---|---|
| AAAAAGCAGGCTTCAACATGAAATTCCTATCACTCGCCTTC | CG16844 primer 1st Fw | BomS3 |
| AGAAAGCTGGGTCCTANTCANTTAGGCCCTCACATTGCAGACG | CG16844 primer 1st RvS | |
| AAAAAGCAGGCTTCAACATGAAGTGGATGTCCTTGGTCT | CG15065 primer 1st Fw | BomS5 |
| AGAAAGCTGGGTCTCANCTANTTAGCCTCCGCGAACATTACAAT | CG15065 primer 1st RvS | |
| AAAAAGCAGGCTTCAACATGAAGAGCCTGACGTTATTG | CG15067 primer 1st Fw | BomBc2 |
| AGAAAGCTGGGTCCTANTCANTTATCCGCGAATATTGCAACCG | CG15067 primer 1st RvS | |
| AAAAAGCAGGCTTCAACATGAAGCTGCTCTCGATTACTTT | CG15068 primer 1st Fw | BomS6 |
| AGAAAGCTGGGTCTTANTCANCTAATCGCCGCGTATATTGCAAA | CG15068 primer 1st RvS | |
| AAAAAGCAGGCTTCAACATGAAAGCTCTTCAAGTCG | CG16836 primer 1st Fw | BomT2 |
| AGAAAGCTGGGTCCTANTCANTTAACTGAATATATGATATATAATTCC | CG16836 primer 1st RvS | |
| AAAAAGCAGGCTTCAACATGAAGTTCTTCTCAGTCGTCA | CG18106 primer 1st Fw | BomS2 |
| AGAAAGCTGGGTCTTANTCANCTACTTTCCACCGTGCACATTG | CG18106 primer 1st RvS | |
| AAAAAGCAGGCTTCAACATGCGATTCTTTGCAATCGTCA | CG18107 primer 1st Fw | BomS4 |
| AGAAAGCTGGGTCTTANTCANCTATTTGCCACCACGAACATTG | CG18107 primer 1st RvS | |
| AAAAAGCAGGCTTCAACATGAAGTGGTTTTCGATTTTGTTTG | CG43202 primer 1st Fw | BomT1 |
| AGAAAGCTGGGTCCTANTCANTTACCATCTTCGAGTAAATTTGATTG | CG43202 primer 1st RvS | |
| AAAAAGCAGGCTTCAAC ATGAAGTGCCTGATTCTGTCCTTTGC | CG15066 primer 1st Fw | BomBc1 |
| AGAAAGCTGGGTCCTANTCANTTAATCCCCGCCCCCGCTGAT | CG15066 primer 1st RvS | |
| AAAAAGCAGGCTTCAACATGAAATTCTTCTCAGTCGTCACCG | CG18108 primer 1st Fw | BomS1 |
| AGAAAGCTGGGTCTCANTTANCTACTTGCCACCGTGGACATT | CG18108 primer 1st RvS | |
| GGGGACAAGTTTGTACAAAAAAGCAGGCT | GATEWAY 2nd-Fw | primer for sencond round PCR to build attB sequence |
| GGGGACCACTTTGTACAAGAAAGCTGGGT | GATEWAY 2nd-Rv | |
| TGGACCTGCGGGGTTAATTTACC | HL1-F | for barcode amplification |
| TTGTGTCATGTCGGCGACCCTACG | attP-R | |
| GCAACTACTGAAATCTGCCAAG | hsp-GW-F | amplify ORF of interesting gene in transgenic fly |
| AATTAACCCCGCAGGTCCACCGG | HL1-R | |

**Table S2: Inserted ORF sequence and the corresponding barcode sequence for each transgenic Bomanin flies.**

| | clone sequence (ORF) | barcode sequence | translation sequence |
|---|---|---|---|
| BomT1-W1118 | ATGAAGTGGTTTTCGATTTTGTTTGCGCTACTGGCGCTCATTTCCTTCGTCGAATTTGGATATGCTCGTAAGTTCTATCTATAAAAATTATTCCAATGAATATCTATAAAAATAT<br>GTTACTTATTTCAGGAACAATTCCCAGGATTACCATTCGCAATGGCGACATTATAGTTCATGGCAATTGCAATGGCTGCACGGCACGCGCCACCAAAAACTCGGCTCACTTATC<br>AATCAAATTTACTCGAAGATGGTAA | GGTGTGTACTTTGTGAGTAGACGAGATAAAGAGA | MKWFSILFALLALI**S**FV<br>EFGYARTIPRITIRNGDI<br>IVHGNCN**G**CTARATKN<br>SAHLSIKFTRRW |
| BomBc1-W1118 | ATGAAGTGCCTGATTCTGTCCTTTGCAATTTTCGTTGTCCTGGCTTCCCAGGCTACGGGTAAGAATCAGATGGGCGCGCCTTTCAATGTCACAATTTCGCTTATTATTTTATTAT<br>TCCGCAGCCGGAAATGTGATTATCGGCGGTGTATGCCAGGATTGCAGTCCGCCGGTGGCGGAAAACGTGGTGGTCGGTGGCCAATCCTACAGGACGGGTAGGCCGGGCCAGG<br>GAACGGTCTATATCAATTCCCCTGGCGCATATCCAGGAGCTCTCGATGGTCCCATTCGGCGAACTGGCGCTGGCGGCGGAGGAGGCGGTGGCACCCGGTTTTCGGATGGCTATA<br>GTGGTCGTCTGCCAGGTGGCACTTACCTTTACAATAAGGATTGCGTGGGCTGCAGCATCAGCGGGGGCGGGGATTAA | ATTGGCAACGCACTGACTGCACGTCATGTGGATT | MKCLILSFAIFVVLASQ<br>ATAGNVIIGGVCQDCSP<br>PVAENVVVGGQSYRTG<br>RPGQGTVYINSPGAY**P**<br>GALDGPIPRTGAGGGG |
| BomBc1-Canton S | ATGAAGTGCCTGATTCTGTCCTTTGCAATTTTCGTTGTCCTGGCTTCCCAGGCTACGGGTAAGAATCAGATGGGCGCGCCTTTCAATGTCACAATTTCGCTTATTATTTTATTAT<br>TCCGCAGCCGGAAATGTGATTATCGGCGGTGTATGCCAGGATTGCAGTCCGCCGGTGGCGGAAAACGTGGTGGTCGGTGGCCAATCCTACAGGACGGGTAGGCCGGGCCAGG<br>GAACGGTCTATATCAATTCCCCTGGCGCATATCCAGGAGCTCTCGATGGTCCCATTCGGCGAACTGGCGCTGGCGGCGGAGGAGGCGGTGGCACCCGGTATCCGGATGGCTAC<br>AGTGGTCGTCTGCCAGGTGGCACTTACCTTCACAATAAGGATTGCGTGGGCTGCAGCATCAGCGGGGGCGGGGATTAA | AATGCAGAC.TACTGACAACACAGCATGTCGACT | MKCLILSFAIFVVLASQ<br>ATAGNVIIGGVCQDCSP<br>PVAENVVVGGQSYRTG<br>RPGQGTVYINSPGAY**P**<br>GALDGPIPRTGAGGGG |
| BomS1 (CG18108) | ATGAAATTCTTCTCAGTCGTCACCGTTTTTGTGCTCGGTCTGCTGGCTGTGGCCAATGGTGAGTAAAACTAATTGAAATTGCCTATTTGATGATTTAACTAATAAATTATTATT<br>TTACAGCTGTTCCACTGTCGCCCTATCCAGGAAATGTGATCATCAATGGCGATTGCAGAGTGTGCAATGTCCACGGTGGCAAGTAG | TATGACGACCCCCTGAATCCACAGGATCTAGATG | MKFFSVVTVFVLGLLA<br>VANAVPLSP**Y**PGNVIIN<br>GDCRVCNVHGGK |
| BomS4 (CG18107) | ATGCGATTCTTTGCAATCGTCACTGTCTTTGTGCTTGGTCTTCTGGCTTTGGCCAATGGTGTGTAGAACCTACAAAGATTATTTAGTTGATTACTTAAGTAAATAATCATCTTAT<br>TCAACAGCTGTTCCGTTGTCACCCGATCCAGGAAATGTTATCATCAATGGCGATTGTGTGAATTGCAATGTTCGTGGTGGCAAATAG | TATGTTGAC.CATCTGACGGTACTGGATGATGAG | KGGRANCTVCDGGIIA<br>NAPAPSLP**V**ANGLALL<br>GGVCVTCIAFGGM |
| BomBc2 (CG15067) | ATGAAGAGCCTGACGTTATTGGCGCTTTTGATCTTGGCCACGTTGGCCTTTGTCCACGGTCGGTTTGAAGGCCAAAAGGTTTTAACTTGAATATTAACCGTTCTTTCTTTCTTCT<br>ACCTAGGTGGCAAGGTGACCATTAACGGCAAGTGCGTAAACTGCTCCCATGATCAAACGACTACCACCACGCACAAGCCAACAAGTGGAAAAGGAAGTGGAGGTAGGACGAC<br>AGCCCGGCCAAGCTCGAGATCATCTCCATCCCGCGGTCGCCCATCATGGGATGACGATGATGATGATGATGACCTGACCGGTGGCTGGTCACTGCATCAATCGGCTGGAGGAA<br>CTCAATACATTGGCAGGCGTTCGAAGCGTCAGTCACGCGGTGGTCAGTACATCGATTTGGGCGGATCTGGCGGAAGGGGTGGAGGCGGAGGCTGGGCAGGATCTGGCATAAC<br>CACCATCGATTCCAGTGGCTATCCCGGCGGAACCCTGGTGCGCAACAGTGATTGCGTCGGTTGCAATATTCGCGGATAA | CCTGTAGACGAGCTGACGTCACGAGATTTTGAGA | MKSLTLLALLILATLAF<br>VHGGKVTINGKCVNCS<br>HDQTTTTTHKPTSGKG<br>SGGRTTARPSSRSSP**S**R<br>GRPSW**D**DDDDDDDLT<br>G**G**W**S**LHQSAGGTQYIG |
| BomS2 (CG18106) | ATGAAGTTCTTCTCAGTCGTCACCGTCTTTGTGTTCGGTCTGCTGGCTCTGGCCAACGGTTAGTAATACTATATTTTATTGCTATTTGATTACATTATTAATTATTTATTTCTTCC<br>GCAGCTGTTCCCCTGTCGCCCGATCCAGGAAATGTGGTAATCAACGGCGACTGCAAATACTGCAATGTGCACGGTGGAAAGTAG | ATTGCGTACCACGTGAGCTGACCACATATAGAAT | MKFFSVVTVFVFGLLA<br>LANAVPLSPDPGNVVI<br>NGDCKYCNVHGGK |
| BomS3 (CG16844) | ATGAAATTCCTATCACTCGCCTTCGTTTTGGGTCTGCTGGCTCTGGCCAACGGTGAGTAATGCCAACTATTTACAAACTAAATAAGTTATTTAATTATGATTTTAATTATATTTC<br>CAACACAGCCACTCCCCTGAATCCTGGCAATGTCATCATCAATGGCGATTGCCGCGTCTGCAATGTGAGGGCCTAA | CCTGCGCACCTGATGAGTATACGTAATATCGAGC | MKFLSLAFVLGLLALA<br>NATPLNPGNVIINGDCR<br>VCNVRA |
| BomT2 (CG16836) | ATGAAAGCTCTTCAAGTCGCCGGAACTTTGATGCTGCTTTTCTGCCTGCTGGCAGCTGTTAATGGTAATTATTTAATGAAATAATAAAATATATATATAATATTTTTAGCTAAT<br>CATTTCCTTATTGCAGCTACGCCGGGACAAGTGTATATCAATGGGAAATGCATTGACTGCAATAAGCCTGATAATGATCCGGGAATTATAATTCCTCCAGACCATAAATCAGCT<br>GGATCCATGTCTTACACACTCACATCTGGAGCCATCTTCTTTGGAATTATATATCATATATTCAGTTAA | CTTGACCACCCTATGAGGGCACAGAATGAAGAGT | MKALQVAGTLMLLFC<br>LLAAVNATPGQVYING<br>KCIDCNKPDNDPGIIIPP<br>DHKSAGSMSYTLTSGA<br>IFFGIIYHIFS |
| BomS5 (CG15065) | ATGAAGTGGATGTCCTTGGTCTTTCTATGCGGTCTGCTCGCCATGGCAGTGGGTGAGTATCTATAAGATTCATTAATCCTTAACCAGTATCCTAATTATTTTTCTACTATTACTA<br>TAGCTTCTCCGTTAAATCCGGGTAATGTCATTATCAATGGAGATTGTCGTCATTGTAATGTTCGCGGAGGCTAA | ACTGACCACCTGATGAGAACACCTGATGTTGATC | MKWMSLVFLCGLLAM<br>AVASPLNPGNVIINGDC<br>RHCNVRGG |
| BomS6 (CG15068) | ATGAAGCTGCTCTCGATTACTTTTCTCTTATGAAGCTGCTCTCGATTACTTTTCTCTTCGGACTTTTGGCTTTGGCTAGTGGTAAGTGCCAATGATGTGCTGTATAAATAATAAA<br>AACCTATTTCTAATTGAAATCTTTATTTTATTTACTTATTTCTAGCCAATCCCCTGAGTCCTGGCAATGTGATTATAAACGGCGACTGCAAAGTTTGCAATATACGCGGCGATTA<br>G | CTTGCTGACGGCCTGAGCAGACGCCATATAGACA | MKLLSITFLFGLLALAS<br>ANPLSPGNVIINGDCKV<br>CNIRGD |

#### Table S3: Primer sequence of the gRNA plasmid construction.

| | |
|---|---|
| GCGGCCCGGGTTCGATTCCCGGCCGATGCATTTCGTTGTCCTGGCTTCCCGTTTTAGAGCTAGAAATAGCAAG | IM23-gSingle-fwd |
| ATTTTAACTTGCTATTTCTAGCTCTAAAACGGTGGCACTTACCTTCACAATGCACCAGCCGGGAATCGAACCC | IM23-gSingle-Rvs |
| GCGGCCCGGGTTCGATTCCCGGCCGATGCATGATCATCAATGGCGATTGCGTTTTAGAGCTAGAAATAGCAAG | IM1-gSingle-fwd |
| ATTTTAACTTGCTATTTCTAGCTCTAAAACGTGCACTCAGTATCCAAAACTGCACCAGCCGGGAATCGAACCC | IM1-gSingle-Rvs |
| GCGGCCCGGGTTCGATTCCCGGCCGATGCAGTGAATTGCAATGTTCGTGGGTTTTAGAGCTAGAAATAGCAAG | CG18107-gSingle-fwd |
| ATTTTAACTTGCTATTTCTAGCTCTAAAACATGgtgtgtagaacctacaaTGCACCAGCCGGGAATCGAACCC | CG18107-gSingle-Rvs |
| GCGGCCCGGGTTCGATTCCCGGCCGATGCATGTTTAAACTTATGTGGTGGGTTTTAGAGCTAGAAATAGCAAG | CG15067-gSingle-fwd |
| ATTTTAACTTGCTATTTCTAGCTCTAAAACGACGTTATTGGCGCTTTTGATGCACCAGCCGGGAATCGAACCC | CG15067-gSingle-Rvs |
| GCGGCCCGGGTTCGATTCCCGGCCGATGCATTGAATTCAACTGATGCTCTGTTTTAGAGCTAGAAATAGCAAG | IM2-gSingle-fwd |
| ATTTTAACTTGCTATTTCTAGCTCTAAAACGTCGCCCGATCCAGGAAATGTGCACCAGCCGGGAATCGAACCC | IM2-gSingle-Rvs |
| GCGGCCCGGGTTCGATTCCCGGCCGATGCAACTCGGGAATTTCTCGATGGGTTTTAGAGCTAGAAATAGCAAG | IM3-gSingle-fwd |
| ATTTTAACTTGCTATTTCTAGCTCTAAAACCGTCTGCAATGTGAGGGCCTTGCACCAGCCGGGAATCGAACCC | IM3-gSingle-Rvs |
| GCGGCCCGGGTTCGATTCCCGGCCGATGCAATGAAAGCTCTTCAAGTCGCGTTTTAGAGCTAGAAATAGCAAG | IM28-gSingle-fwd |
| ATTTTAACTTGCTATTTCTAGCTCTAAAACGCTGATTTATGGTCTGGAGGTGCACCAGCCGGGAATCGAACCC | IM28-gSingle-Rvs |
| GCGGCCCGGGTTCGATTCCCGGCCGATGCATCTATGCGGTCTGCTCGCCAGTTTTAGAGCTAGAAATAGCAAG | CG15065-gSingle-fwd |
| ATTTTAACTTGCTATTTCTAGCTCTAAAACCGAACATTACAATGACGGCATGCACCAGCCGGGAATCGAACCC | CG15065-gSingle-Rvs |
| GCGGCCCGGGTTCGATTCCCGGCCGATGCACTAGCCAATCCCCTGAGTCCGTTTTAGAGCTAGAAATAGCAAG | CG15068-gSingle-fwd |
| ATTTTAACTTGCTATTTCTAGCTCTAAAACctggagtatataaaaatgcgTGCACCAGCCGGGAATCGAACCC | CG15068-gSingle-Rvs |

**Table S4: Primers to identify knock-in/out mutant flies for BomS2, BomT2, and BomS5.**

| | Transgene-specific primer | Sequence |
|---|---|---|
| BomS2 | IM2KI-6-F | TGCTGCTCCAAGATTCCAGG |
| | IM2KI-6-R | AGCTGCAAGGTATTAAAAATACGAA |
| BomT2 | CG16836KI-8-F | TCAGCCTTGCTACTGTATGGT |
| | CG16836KI-8-R | AAGTTCGCAATTTGTCACAAGC |
| BomS5 | CG15065-F | AATCAAACCGATTTGCGCGG |
| | CG15065-R | CTTGAAGCTGTCCTTCCCCG |

**Table S5: Primers used in quantitative RT-PCR.**

| RT-qPCR primer | primer name |
|---|---|
| CAATGCTGTTCCACTGTCGC | BomS1-Fq |
| CGTGGACATTGCACACCCTG | BomS1-Rq |
| TTTCGTTGTCCTGGCTTCCC | BomBc1-Fq |
| CCGACTACGACGTTTTCCGC | BomBc1-Rq |
| AGAGCCTGACGTTATTGGCG | BomBc2-Fq |
| TGGGAGCAGTTTACGCACTT | BomBc2-Rq |
| AGTCGTCACCGTCTTTGTGTT | BomS2-Fq |
| CAGTATTTGCAGTCCCCGTTG | BomS2-Rq |
| TCACTCGCCTTCGTTTTGGG | BomS3-Fq |
| TTAGGCCCTCACATTGCAGAC | BomS3-Rq |
| TAATGCTACGCCGGGACAAG | BomT2-Fq |
| ATGGCTCCAGATGTGAGTGTG | BomT2-Rq |
| GTTGTCACCCGATCCAGGAA | BomS4-Fq |
| ATTTGCCACCACGAACATTGC | BomS4-Rq |
| TCCCAGGATTACCATTCGCA | BomT1-Fq |
| GAGTTTTTGGTGGCGCGTG | BomT1-Rq |
| GGTCTTTCTATGCGGTCTGCT | BomS5-Fq |
| TAGCCTCCGCGAACATTACA | BomS5-Rq |
| GCTAGTGCCAATCCCCTGAG | BomS6-Fq |
| TCGCCGCGTATATTGCAAAC | BomS6-Rq |
